# Sociality shapes mitochondrial adaptations supporting hypoxia tolerance

**DOI:** 10.1101/2024.09.30.615914

**Authors:** Alice Rossi, Max Ruwolt, Paraskevi Kakouri, Tetiana Kosten, Severine Kunz, Dmytro Puchkov, Jane Reznick, Bettina Purfürst, Damir Omerbašić, Daniel Méndez Aranda, Giorgia Carai, Guido Mastrobuoni, Daniel W. Hart, Michela Carraro, Ludovica Tommasin, Nigel C. Bennett, Valérie Bégay, Katja Faelber, Oliver Daumke, Paolo Bernardi, Thomas J. Park, Stefan Kempa, Fan Liu, Gary R. Lewin

## Abstract

Oxygen deprivation or hypoxia is poorly dealt with by most terrestrial species and often leads to permanent tissue damage and death. One prominent exception is the naked mole-rat (*Heterocephalus glaber*) which is remarkably adapted to withstand prolonged periods (∼18 mins) of severe hypoxia, a trait likely driven by its crowded underground lifestyle. Other African mole-rat species are less social or entirely solitary like the Cape mole-rat (*Georychus capensis*). Here, we asked whether cellular and molecular adaptations to hypoxia map to social traits. We discovered that at the cellular level naked mole-rat fibroblasts survive >30 hours in 1% oxygen, while fibroblasts from terrestrial or non-social mole-rat species (human, mouse and Cape mole-rat) die rapidly under hypoxic conditions. We further show that naked mole-rat mitochondria have evolved morphological, functional and proteomic adaptations crucial for hypoxia resistance, remaining unaffected after long periods of severe hypoxia. We identify the mitochondrial protein Optic Atrophy 1 (OPA1) as a key player in naked mole-rat hypoxia resilience. Naked mole-rat mitochondria not only express more protective forms of OPA1, but also harbor a structurally unique isoform that likely protects cells from hypoxic damage. We show that evolutionary changes including the functionalization of a unique *Opa1* exon support mitochondrial mediated cellular protection. Indeed, knockdown of OPA1 in naked mole-rat cells is sufficient to render them equally susceptible to hypoxia as human cells or cells from non-social African species. Our study demonstrates how molecular evolution drives unique adaptations that enable cells to achieve unprecedented resistance to hypoxic damage. We also show that molecular changes at the level of mitochondria are crucial in conferring hypoxia resistance. Our results thus chart a novel molecular path to understand how robust cellular hypoxia resistance can be achieved. Such knowledge may eventually inspire novel strategies to circumvent the consequences of hypoxic-damage in humans.

## Introduction

Oxygen is essential for most invertebrate and vertebrate life on Earth. Short or prolonged periods of oxygen deprivation in mammals due to ischemic episodes lead to irreversible tissue damage and often death in most species^1^. However, some exceptional vertebrate species including pond turtles, crucian carp and snakes have evolved mechanisms to live with extremely low oxygen availability that would be lethal to most mammals^2,3^. However, one mammalian species, the naked mole-rat (*Heterocephalus glaber),* has evolved exceptional hypoxia resistance^4–6^. This eusocial species is adapted to live in large underground groups and is thought to be exposed to regular bouts of hypoxia and hypercapnia^7^. Hypoxic conditions for naked mole-rats may be especially serious during sleep, as a signature behavior of this subterranean mammal is that they sleep communally occupying self-built chambers, barely large enough to accommodate all animals in the group^6^. Therefore, it is unsurprising that naked mole-rats can tolerate long periods of both hypoxia (5% O_2_) and anoxia (0% O_2_) without any organ damage^4^. Naked mole-rats belong to the *Bathyergidae* family, which comprises more than 30 fossorial African mole-rat species and occupy the full range of the sociality spectrum from eusocial (animals live in large colonies of up to 300 individuals), to social (animals live in small family groups) to solitary^8^. We hypothesized that the ability to survive hypoxia scales with sociality within the *Bathyergidae* family, where animals that live in large colonies like naked mole-rats may have acquired unique physiological mechanisms to better cope with the low amount of oxygen available in small and crowded burrows^6^. Several studies have indicated that naked mole-rat tissues have adapted to hypoxic stresses through changes in gene expression that for example may promote altered mitochondrial function^9–11^. Furthermore, mitochondria are prime targets for hypoxic stress and normally initiate cell death after fragmentation^12,13^. To examine more directly whether naked mole-rat mitochondria are adapted to hypoxia, we established primary fibroblasts as a model system. Here we could also compare cellular hypoxia resistance between cells from the naked mole-at and multiple hypoxia-prone species. These models enabled us to pinpoint morphological, functional and proteome adaptions of naked mole-rat mitochondria that are required to protect cells from prolonged hypoxia. Our comparative and cell biological analyses allowed us to identify genome changes that may lead to alterations in the oligomeric structure of the dynamin-like GTPase OPA1, a fusion and cristae modelling protein, that hinders cell death initiation in naked mole-rat.

## Results

### Anoxia resistance scales with sociality

We used an extreme anoxic challenge to compare in vivo anoxia resistance across members of the *Bathyergidae* family^4^. Mice (*Mus Musculus*), rats (*Rattus norvegicus*) and four African mole-rat species with different social structures and group sizes - Damaraland (*Fukomys damarensis*; eusocial), Natal (*Cryptomys hottentotus natalensis*; social), Mahali (*Cryptomys hottentotus mahali*; social) and Cape (*Georychus capensis*; solitary) were challenged^8^ (Fig. 1a). Animals were exposed to 0% O_2_ by using a chamber flushed with N_2_ (10 l/min), all species ceased voluntary movement in less than 50 seconds and were clearly unconscious. However, the two eusocial species, naked and Damaraland mole-rats, continued to make breathing attempts for several minutes^4^ (Fig. 1b). The social species, Natal and Mahali mole-rats, made breathing attempts for less than 2 minutes, and the solitary Cape mole-rat, rat and mouse for less than 1 minute (Fig. 1b). In this protocol animals were re-exposed to room air one minute after the last breathing attempt. Despite spending several minutes exposed to anoxia all naked mole-rats revived in room air (Fig. 1b). Half of the Damaraland mole-rats also revived completely after exposure to room air, but none of the social Mahali mole-rats survived the exposure. The social Natal mole-rat showed robust recovery but was exposed to much shorter periods of anoxia than the eusocial species. Notably, the solitary Cape mole-rat did not generally recover from anoxia and thus showed a similar vulnerability to anoxia as non-fossorial rodents like mice and rats (Fig. 1b). Thus, hypoxia resistance appears to scale with sociality and is not a general feature of all African mole-rats.

**Figure 1.**
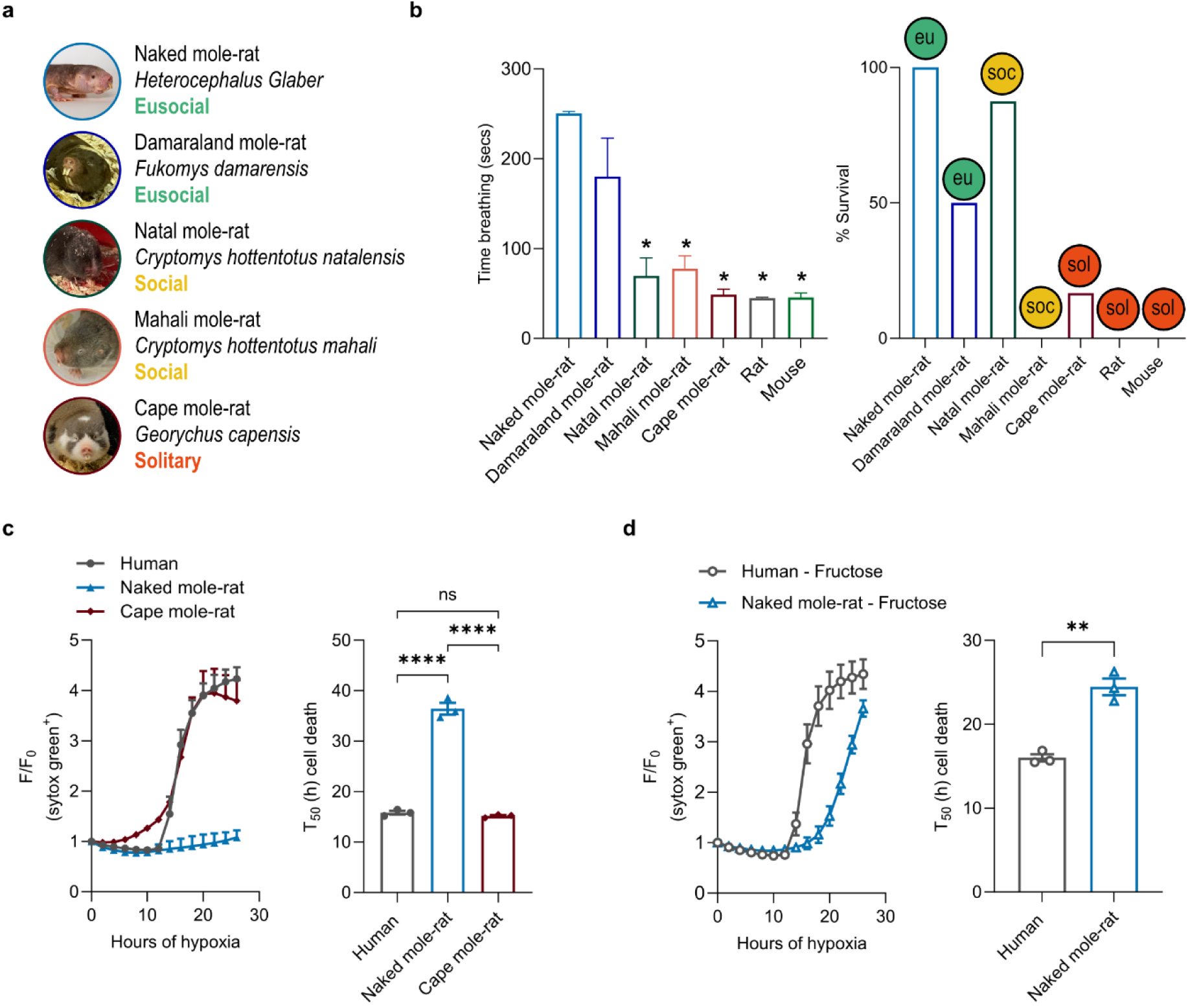
Hypoxia resistance scales with sociality in mammalian species. **(a)** Five eusocial (eu), social (soc) and solitary (sol) African mole-rats species belonging to the *Bathyergidae* family studied in the anoxia resistance experiment. **(b)** The mean time breathing (left) and the percentage of survival (right) were calculated upon exposure to anoxia (0% O_2_) in mouse, rat and in the five African mole-rat species. n= number of animals, n ≥ 4. One-way ANOVA followed by multiple comparison test to naked mole-rat. The naked mole-rat and mouse data are from^4^. **(c)** Mean survival curve (left) and cell death time 50 (T_50_) (right) in human, naked mole-rat and cape mole-rat primary fibroblasts exposed to 24 h of hypoxia (1% O_2_). **(d)** Mean survival curve (left) and cell death time 50 (T_50_) (right) in human and naked mole-rat primary fibroblasts cultured in medium deprived of glucose and supplemented of fructose (see Methods) and exposed to 24 h of hypoxia (1% O_2_). (c-d) Each dot (n) is the number of experiments, n=3. One-way ANOVA (c) and Student’s t test (d). *p < 0.05, **p < 0.01, ****p < 0.0001. Data are presented as mean values ± s.e.m.

We next asked whether naked mole-rat cells are intrinsically hypoxia-resistant. We used primary fibroblasts as a model which enabled us to compare the hypoxia susceptibility, not only of naked mole-rat, mouse and human cells, but also of cells from the solitary Cape mole-rat (Extended Data Fig. 1a). When exposed to 1% oxygen, fibroblasts from human, mouse and the Cape mole-rat showed a rapid decline in cell viability with 50% of the cells dead 16 h later (mean T_50_ ∼ 16 h; Fig. 1c and Extended Data Fig. 1, a-c). In sharp contrast, naked mole-rat fibroblasts showed a remarkable ability to cope with such extreme conditions with T_50_ > 35 h (Fig. 1c; Extended Data Fig. 1d), and many cells remained viable up to 48 h after hypoxia exposure (Extended Data Fig. 1d). Most of our experiments were done with neonatal kidney-derived fibroblasts, however, we noted similar hypoxia resilience in fibroblasts from the skin and the kidney taken from older animals (Extended Data Fig. 1 a, e). We previously showed that naked mole-rat tissues can switch to fructolysis under hypoxic conditions^4^ and asked if this is also the case for naked mole-rat fibroblasts. When we exposed naked mole-rat fibroblasts to hypoxia in media where glucose had been replaced with fructose we again observed that naked mole-rat fibroblasts survived very well with T_50_ > 20 h, in contrast, human fibroblasts died rapidly (Fig. 1d). Taken together, these data indicate that naked mole-rat fibroblasts exhibit remarkable cellular adaptation to extreme hypoxia, not seen in other mammalian cells, including fibroblasts obtained from a related hypoxia susceptible African mole-rat species.

### The naked mole-rat proteome reveals a unique mitochondrial biology

To uncover the molecular underpinnings underlying the differences between the hypoxia-resistant naked mole-rat and the mouse, we used label-free proteomics to compare protein abundances in the liver, a high energy demand tissue. Orthologous peptides were identified in parallel Mass Spectrometry runs allowing us to make a cross-species protein identification using mouse databases^14–16^. We obtained robust measurements of 1313 liver proteins. Compared to the mouse, 48% of naked mole-rat proteins showed abundances that were at least two-fold different between the two species. Using a gene ontology analysis (GO) of cellular components and Kyoto Encyclopedia of Genes and Genomes (KEGG pathways) we showed a predominant downregulation of proteins localized in mitochondria and associated with metabolic pathways (Fig.2, a, b), while glycolysis, fructose metabolism and proteasome pathways were upregulated (Extended Data Fig. 2 a, b). This analysis was consistent with previous studies^4,10^ and prompted us to investigate in more detail mechanistic links between mitochondria and hypoxia resistance. First, we examined the fine structure of liver mitochondria using Transmission Electron Microscopy (TEM) with the cryo-sectioning method of Tokuyasu which gives an excellent resolution of mitochondrial membranes^17^ (Fig. 2c-e). Analysis showed that liver mitochondria from the naked mole-rat were smaller than those of mice (Fig. 2c, d), but more strikingly exhibited a marked scarcity of cristae (Fig. 2c, e). Mitochondrial morphology often correlates with their functionality^18^, and we, therefore, performed oxygen consumption measurements in isolated mitochondria from mouse and naked mole-rat liver (Fig. 2f-h and Extended Data Fig. 2c, d). Respiration rates were evaluated using glutamate/malate (G/M) as substrates for complex I (Fig. 2g) or succinate/rotenone (Succ/Rot) as substrates for complex II and inhibitor of complex I, respectively (Fig. 2h). Following the addition of ADP (ATP synthase substrate), an increase in respiration rate, then blocked by oligomycin (ATP synthase blocker) was observed (Fig. 2g, h and Extended Data Fig. 2c). Naked mole-rat mitochondria showed lower oxygen consumption rates when supplemented with Succ/Rot compared to mice (Fig. 2g), suggesting reduced complex II activity. Upon ADP addition, naked mole-rat mitochondria also showed lower respiration rates compared to mice, regardless of the substrates (Fig. 2g, h). The latter finding suggests that there is a relative inability to utilize ADP for ATP synthesis. Consistent with whole tissue measurements there was a significantly reduced maximal respiration capacity of naked mole-rat mitochondria, as measured after the addition of a mitochondrial oxidative phosphorylation uncoupler, FCCP (Fig. 2g, h and Extended Data Fig. 2 c, d)^10^. Besides the reduced mitochondrial activity observed in naked mole-rat, the analysis of the ATP synthase organization in isolated mitochondria from mice and naked mole-rats liver showed no dramatic differences between the two species (Extended Data Fig. 2 e, f). These findings indicate that naked mole-rat tissues harbor mitochondria with morphologies consistent with altered function, including reduced mitochondrial respiratory capacity. The mitochondrial adaptations we observed in naked mole-rat are likely relevant for cellular hypoxia resistance.

**Figure 2.**
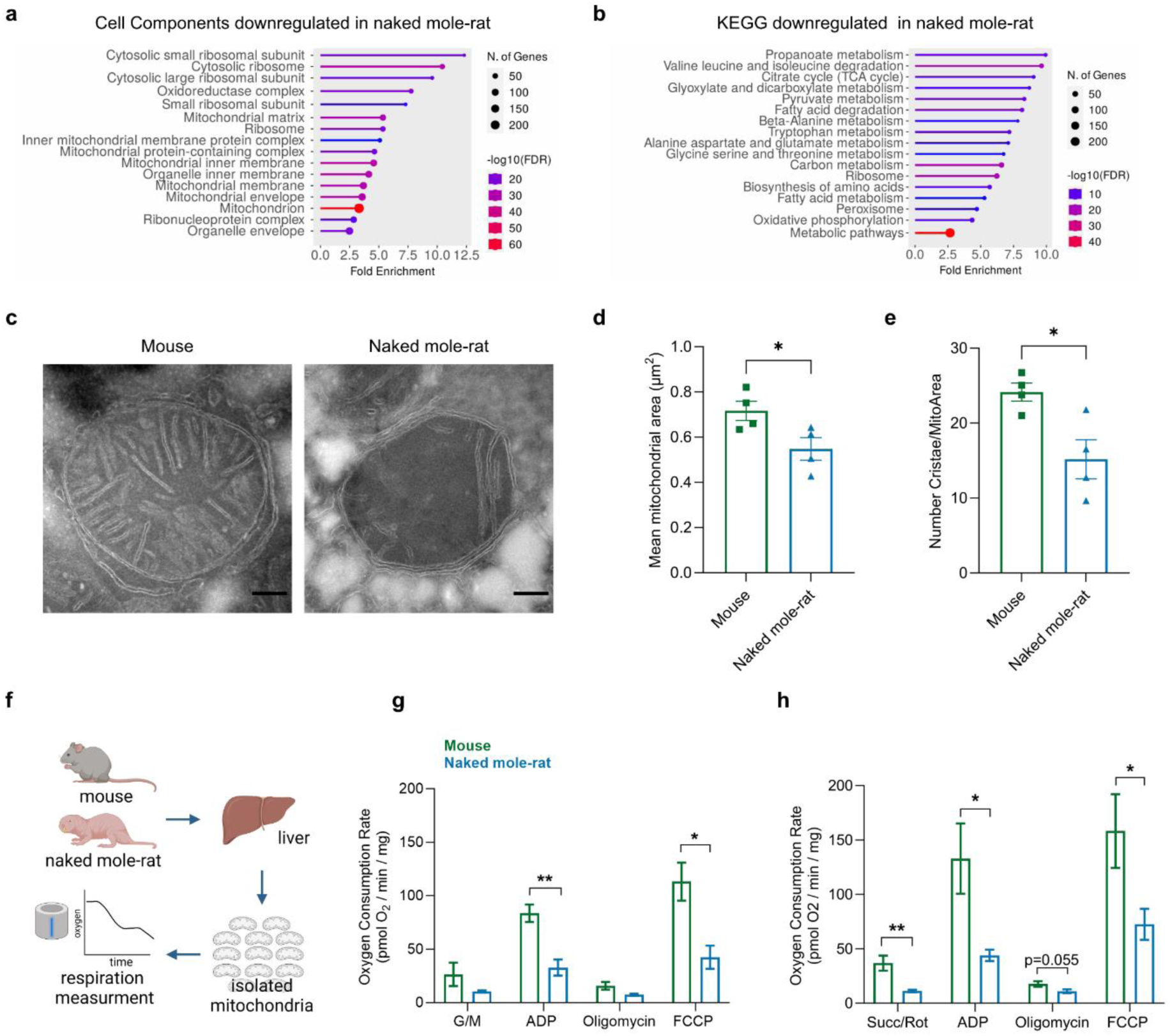
Naked mole-rat liver mitochondria show altered morphology and function in normoxia (a-b) Cell components and KEGG Gene Ontology (GO) term enrichment analysis of naked mole-rat downregulated proteins compared to mouse. **(c-d-e)** Transmission Electron Microscopy analysis of mouse and naked mole-rat liver mitochondria. **(c)** Representative pictures of mouse (left) and naked mole-rat (right) mitochondria. Scale bar 200nm. **(d-e)** Mitochondrial morphology analysis: mean mitochondrial area and number of cristae/mitochondrial area are analyzed. Student’s t-test. Each dot (n) is the number of animals, n≥3. **(f-g-h)** Oxygen Consumption Rate analysis in mouse and naked mole-rat isolated mitochondria. **(f)** Representative scheme of the mitochondria isolation and of the oxygen consumption measurements. **(g-h)** Oxygen consumption Rate when Glutamate/Malate (G/M; **h**) or Succinate/Rotenone (Succ/Rot; I) are provided as mitochondrial substrates. n is number of animals, n=3. Multiple t-test. *p < 0.05, **p < 0.01. Data are presented as mean values ± s.e.m.

### Naked mole-rat mitochondria are key to hypoxia resilience

Fibroblasts from the eusocial naked mole-rat, but not from the solitary Cape mole-rat, humans or mice, show remarkable hypoxia resistance. We next asked whether the mitochondria of hypoxia-resistant fibroblasts showed similar morphologies to those of intact liver. We carried out a quantitative ultrastructural analysis of human and naked mole-rat fibroblasts using transmission electron microscopy (TEM) as well as imaging 3D volumes with focused ion beam scanning electron microscopy (FIB-SEM). Both these methods revealed dramatic differences in mitochondrial morphology between human and naked mole-rat. As in the intact liver, normoxic fibroblast mitochondria were smaller and showed a marked paucity of cristae compared to human cells (Fig. 3a, b and Extended Data Fig. 3a, b). We also looked at mitochondrial ultrastructure 4 hours after oxygen deprivation, and as expected in human cells we observed marked fragmentation reflected by a decrease in mitochondrial perimeter and area, and reduced cristae compared to normoxia^19–21^ (Fig. 3a, b and Extended Data Fig. 3a, b). In contrast, the morphology and size of naked mole-rat mitochondria appeared to be largely unaffected by the lack of oxygen (Fig. 3a, b and Extended Data Fig. 3a, b). We further validated these findings using light microscopy in which we imaged mitochondria using a mitochondrial-targeted GFP delivered via lentiviral constructs or TOM20 labelling followed by confocal imaging (Fig. 3c, d Extended Data Fig. 3 c-g). In line with our TEM analysis (Fig. 3a, b and Extended Data Fig. 3 a, b), under normoxic conditions, the eusocial naked mole-rat fibroblasts exhibited a higher number and smaller mitochondria compared to human and solitary Cape mole-rat cells (Fig. 3c, d and Extended Data Fig. 3c, d). However, upon oxygen deprivation, human and Cape mole-rat mitochondria underwent fragmentation, in sharp contrast naked mole-rat mitochondria remained surprisingly unaffected even after 24 hours of hypoxia (Fig. 3c, d and Extended Data Fig. 3 c, e-g). Similar data were observed in naked mole-rat fibroblasts derived from skin, which also showed no apparent mitochondrial fragmentation in the absence of oxygen (Extended Data Fig. 3h). Taken together, our data indicate that naked mole-rat mitochondria are adapted to low oxygen conditions and do not undergo the dramatic morphological remodeling observed in human and Cape mole-rat fibroblasts during hypoxia. Our data on liver suggests that naked mole-rat predominantly use glycolytic flux to maintain energy homeostasis largely avoiding oxidative respiration in the mitochondria. To test the idea that a relative mitochondria quiescence is necessary for hypoxia resistance we forced our cells to utilize oxidative phosphorylation for ATP production using a medium devoid of glucose and supplemented with galactose and pyruvate^22,23^. Upon adaptation to galactose, we now observed that both naked mole-rat and human cells underwent rapid hypoxia-dependent cell death with kinetics that was virtually identical (Fig. 3e). Furthermore, under galactose media conditions we now observed that both human and naked mole-rat mitochondria underwent fragmentation characterized by decreased mean mitochondrial area and perimeter (Fig. 3f and Extended Data Fig. 4a). Thus, even though under basal conditions naked mole-rat mitochondria are small this does not mean that they cannot undergo fragmentation associated with cell death. During hypoxia, following mitochondrial fragmentation, mitochondrial ATP production is reduced due to the lack of available oxygen^24,25^. Therefore, we measured ATP levels both in glucose and galactose media under normoxic and hypoxic conditions using a bioluminescent assay. Although no detectable differences were observed in the levels of ATP in human and naked mole-rat cells when grown in the two different media (Extended Data Fig. 4b), we found that the level of ATP in naked mole-rat fibroblasts was almost two times higher than in human cells in glucose-containing media (Fig. 3g). As expected, we observed an almost 50% reduction in ATP levels in human fibroblasts upon hypoxia, reflecting the impact of oxygen deprivation (Fig. 3h). In contrast, even with a lower mitochondrial membrane potential (Fig. 3i), naked mole-rat fibroblasts demonstrated a remarkable ability to maintain ATP levels that were not statistically different from those measured under normoxic conditions (Fig. 3h), indicating the preservation of cellular functionality even during prolonged periods of oxygen scarcity. When cells were grown in a galactose medium, we observed that normoxic naked mole-rat fibroblasts were still able to produce a slightly higher amount of ATP compared to human cells, but this difference was not statistically significant (Fig. 3g). In contrast, we found lower ATP levels upon oxygen deprivation similar to those in human cells (Fig. 3j). These findings suggest that naked mole-rat cells, with small mitochondria, low cristae density and depolarized membrane potential, can maintain higher ATP levels compared to human cells both in normoxic and hypoxic conditions when glucose is the energy source. Nevertheless, when mitochondrial activity is forced by the presence of galactose, hypoxia provokes a marked reduction in ATP levels in both species. These results indicate that naked mole-rat cells, like human cells, are susceptible to acute oxygen deprivation, but molecular changes at the level of the mitochondria protect naked mole-rat cells from apoptosis initiation and cell death.

**Figure 3.**
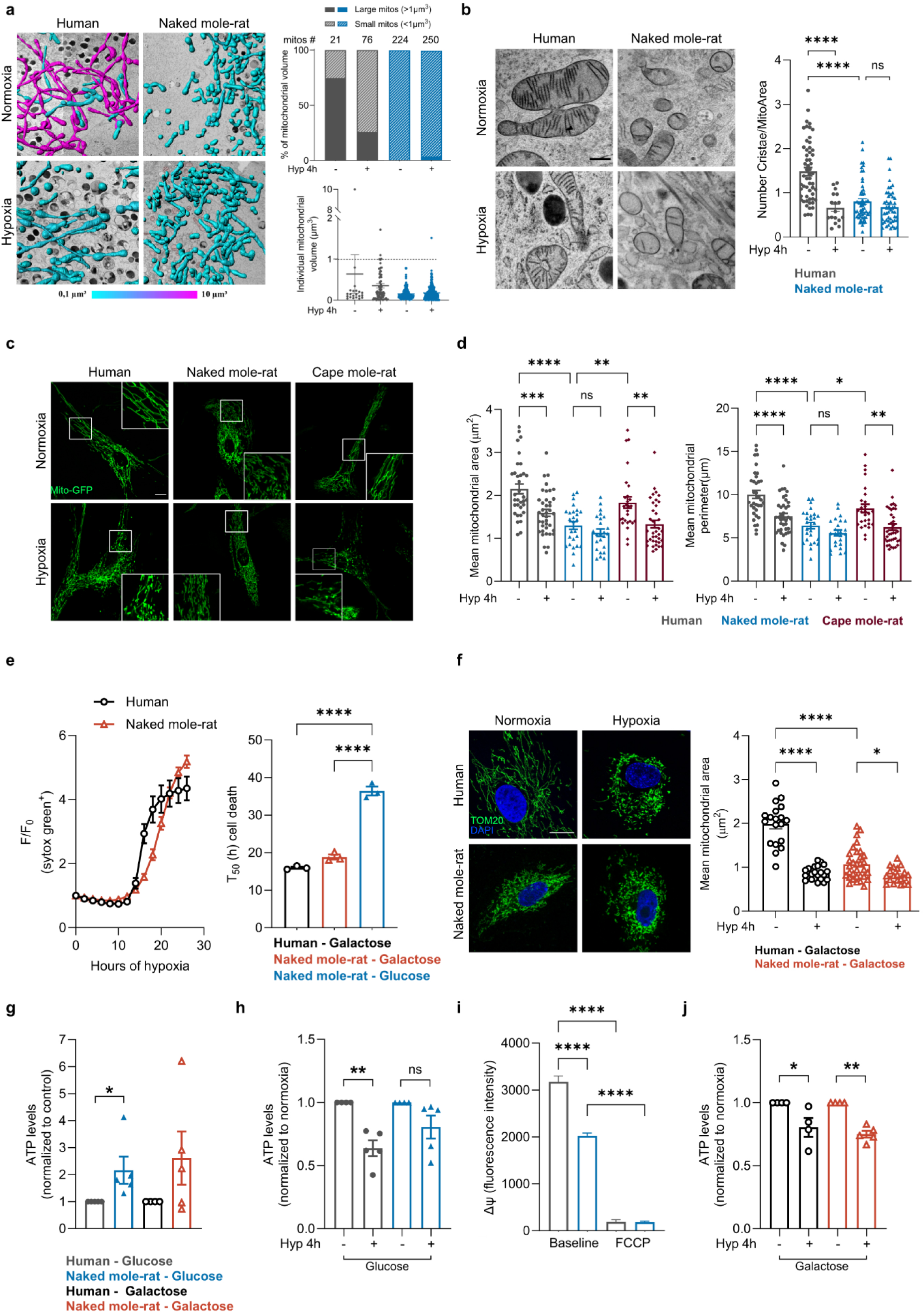
Naked mole-rat mitochondria are resistant to oxygen deprivation. **(a)** 3D reconstruction of mitochondria by FIB-SEM in human and naked mole-rat fibroblasts both in normoxia and after 4 h of hypoxia (1%O_2_). Top right, % of mitochondrial volume occupied by large (>1µm^3^) or small (<1µm^3^) mitochondria. Bottom right, individual mitochondrial volumes. Mitos # is the number of mitochondria. **(b)** Transmission Electron Microscopy analysis of human and naked mole-rat mitochondrial cristae in normoxia and hypoxia (1%O_2_). Left, representative pictures. Scale bar 500nm. Right, quantification of the number of cristae/mitochondrial area. Each dot (n) is number of mitochondria analyzed, n>20 from 3 independent experiments. Normality test followed by Kruskal-Wallis test. **(c-d)** Mitochondrial morphology analysis in human, naked mole-rat and cape mole-rat fibroblasts transduced with mitochondrial-GFP lentivirus, in normoxia and after four hours of hypoxia (1%O_2_). Representative pictures (**c**) and quantification of the mean mitochondrial area (left) and perimeter (right) (**d**) of human, naked mole-rat and cape mole-rat mitochondria. Scale bar 10µm. Each dot (n) is number of cells, n>24 from 3 independent experiments. One-way ANOVA **(e)** Mean survival curve (left) and cell death time 50 (T_50_) (right) in human and naked mole-rat primary fibroblasts cultured in medium deprived of glucose and supplemented with galactose and pyruvate (see Methods) and exposed to 4 h of hypoxia (1%O_2_). The naked mole-rat-glucose bar (light blue) data are the same shown in Fig.1. Each dot (n) represents number of experiments, n=3. One-way ANOVA. **(f)** Mitochondrial morphology analysis of human and naked mole-rat fibroblasts grown in galactose and pyruvate supplemented medium in normoxia and after 4 h of hypoxia (1%O_2_). Left, representative pictures; anti-TOM20 is used to stain mitochondria (green) and Dapi for nuclei (blue). Scale bar 10µm. Right, mean mitochondrial area quantification. n is number of cells; n>15 from 3 independent experiments. One-way ANOVA. **(g-h-j)** Total cellular ATP levels measured in human and naked mole-rat fibroblasts both in glucose and galactose medium in normoxia and upon 4 hours of hypoxia (1%O_2_). (**g**) Data were normalized to human in glucose or galactose medium. (**h-k**). Data were normalized to human or naked mole-rat in normoxic conditions. n is number of experiments, n≥4. One-sample t-test. Data in normoxia glucose and galactose are the same in **g, h, j** and in Extendend Data Fig. 4b. **(i)** Mitochondrial membrane potential (ΔΨ) measurement in human and naked mole-rat fibroblasts by Tetramethylrhodamin-methylester (TMRM). Basal and minimum (induced by FCCP, 10µM) ΔΨ were analyzed. n is number of cells, n>30 from 3 independent experiments. One-way ANOVA. *p < 0.05, **p < 0.01, ***p < 0.001, ****p < 0.0001. Data are presented as mean values ± s.e.m.

### The naked mole-rat mitochondrial proteome is robust in the face of hypoxia

In order to uncover the molecular adaptations enabling naked mole-rat mitochondria to cope with extreme hypoxic conditions, we used TMT-labelling^26^ to specifically examine mitochondrial proteomes. We used biochemical methods to obtain mitochondria-enriched fractions from both human and naked mole-rat fibroblasts under normoxia and hypoxia (Fig. 4a) and identified and quantified 7650 proteins across all four conditions (Extended Data Fig. 4c). Following four hours of exposure to oxygen deprivation, principal component analysis (PCA) consistently depicted a clear separation between human and naked mole-rat-derived mitochondria (Fig. 4b). Notably, substantial proteome remodelling was evident exclusively in human cells, with barely discernible changes observed in naked mole-rat mitochondria between normoxic and hypoxic conditions (Fig. 4b).

**Figure 4.**
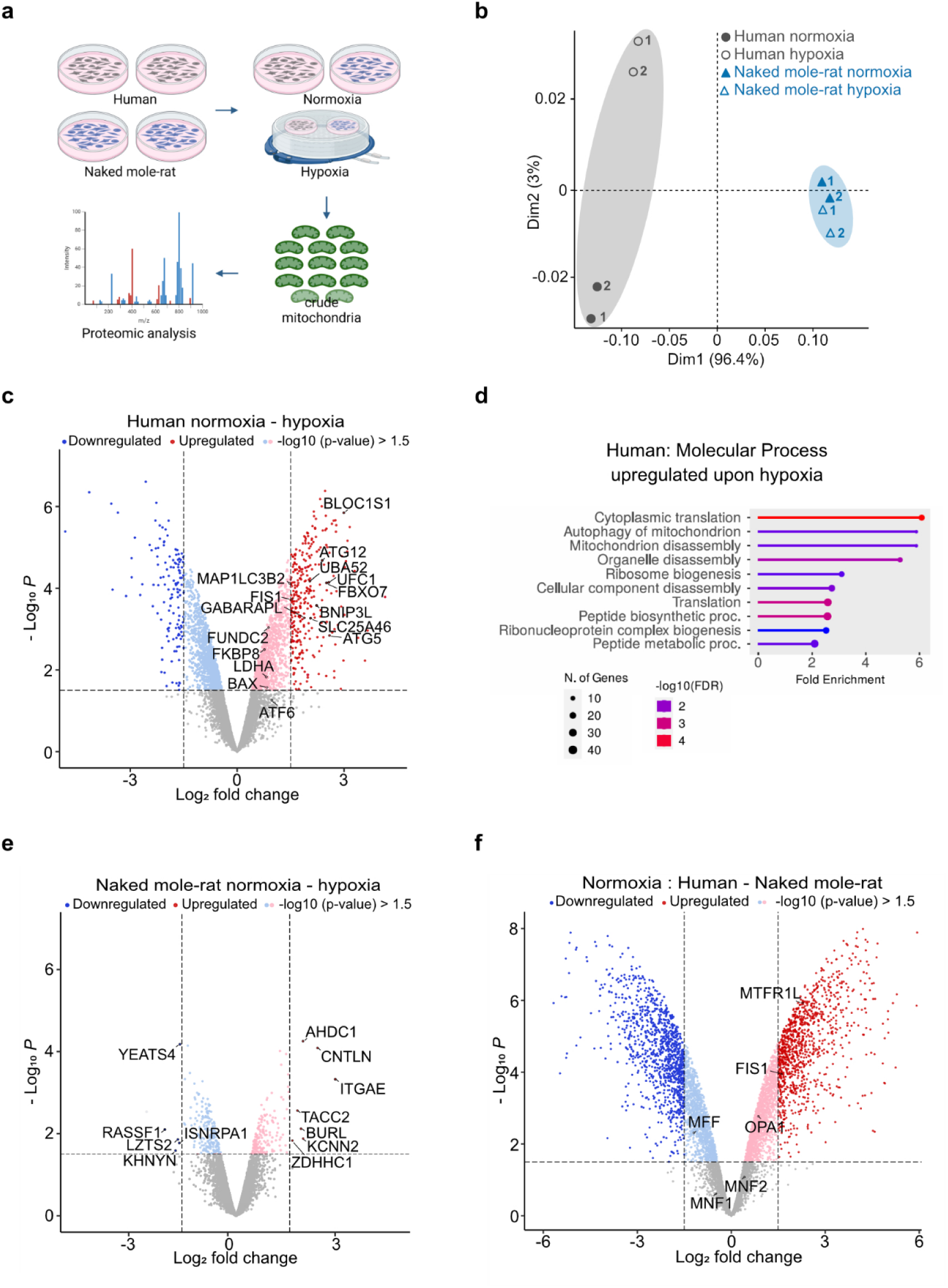
Naked mole-rat mitochondrial proteome does not change upon hypoxia. **(a)** Representative scheme of the proteomic analysis performed in mitochondrial enriched fraction from human and naked mole-rat cells in normoxic conditions and upon 4 h of hypoxia (1%O_2_). **(b)** Principal Component Analysis (PCA) of human and naked mole-rat mitochondrial enriched proteome both in normoxia and hypoxia (1%O_2_). Each dot represents one replicate. **(c)** Volcano plot of up- and downregulated proteins in human fibroblasts upon hypoxia compared to normoxia. Proteins involved in stress response of mitochondria are labelled. **(d)** Molecular Process Gene Ontology (GO) term enrichment analysis of human upregulated proteins upon hypoxia. **(e)** Volcano plot of up- and downregulated proteins in naked mole-rat fibroblasts upon hypoxia compared to normoxia. The 12 up- and downregulated proteins are labelled. **(f)** Volcano plot of up- and downregulated proteins in naked mole-rat fibroblasts compared to human in normoxic conditions. Fission and fusion proteins are labelled. (**c-e-f**) 1.5 Log_2_ fold change and −Log_10_(p-value) cutoff was used.

Using a GO enrichment analysis of cellular component categories we found molecular pathways associated with stress response, mitophagy, organelles degradation and translation activation were upregulated in hypoxic human mitochondria, consistent with previous findings (Fig. 4c, d and Extended Data Fig. 4d) ^25,27,28^. In sharp contrast, the very limited number of dis-regulated proteins (just 12) observed in the naked mole-rat mitochondrial fraction did not show any obvious connection to stress response pathways associated with hypoxic damage (Fig. 4e). In the same experiment, we conducted a cross-species comparative analysis of the mitochondrial proteome between human and naked mole-rat cells under normoxic conditions (Fig. 4f). Interestingly, approximately 50% of the proteins upregulated in response to hypoxia in human cells were already upregulated in naked mole-rat cells under normoxia, suggesting that naked mole-rat mitochondria are pre-adapted to low oxygen.

### OPA1 has naked mole-rat unique structural features required for hypoxia resistance

Given the ultrastructural and metabolic characteristics of naked mole-rat mitochondria observed in normoxia (Fig. 3 and Extended Data Fig. 3), we decided to focus further studies on mitochondrial fission and fusion proteins that might be differentially regulated between the two species (Fig. 4f). Consistent with the predominance of small mitochondria in naked mole-rat cells, we noted upregulation of fission-related proteins, mitochondrial fission protein 1 (Fis1) and the mitochondrial fission regulator 1 like (MTFR1L). We were intrigued to see an up-regulation of the mitochondrial dynamin-like GTPase, Optic Atrophy 1 (OPA1), an important fusion and cristae modelling protein which when deleted in cells leads to fragmented mitochondria with very few cristae (Fig. 4f)^29–33^. Transcriptomic data allowed us to identify two OPA1 C-terminal variants, one similar to the human and Cape mole-rat and one exhibiting an additional stop codon-containing exon (Fig. 5a). Interestingly, the unique naked mole-rat OPA1 isoform was not found in any of the other African mole-rat species, nor any other vertebrate species (Extended Data Fig. 5a). The existence of the additional C-terminal exon in naked mole-rat OPA1 was validated both in fibroblasts and in tissues using RT-PCR (Extended Data Fig. 5b-d). Moreover, homologous DNA sequences in the same genomic location were found in related species and out groups, but these sequences were only functionalized as a coding exon in the naked mole-rat genome (Extended Data Fig. 5e). To explore whether the additional residues at the C-terminal end of naked mole-rat OPA1 may have a functional consequence, we predicted the naked mole-rat OPA1 structure with AlphaFold3 (AF3, Extended Data Fig. 5f) ^34^ and compared it to an experimentally determined cryo-EM structure of oligomerized human OPA1 (Fig. 5b)^35^. Overall, the domain composition and conformation of the G domain, bundle signaling element (BSE), stalk and paddle were highly similar in the two structures (Fig. 5b). Deviations can be found in the third helix of the BSE, which carries a ten amino acid insertion (yellow sequence; Fig. 5 a-c) in naked mole-rat OPA1 and is extended at the C-terminus compared to human OPA1 (green sequence; Fig. 5 a-c). Within the membrane-bound human OPA1 oligomer, the insert might stabilize stalk interface 1 which is essential for oligomer formation^35,36^. Furthermore, the BSE helix extensions of adjacent molecules approach each other and may create a contact, which could also strengthen OPA1 oligomer stability (Fig 5c). Western blot analysis of the long and the short form of OPA1 (L- and S- OPA1) corroborated the upregulation of OPA1 in naked mole-rat, validating our proteomic analysis (Fig. 5d and Extended Data Fig. 6a). We observed a preponderance of S- OPA1 (circa three-fold) compared to the human and Cape mole-rat (Extended Data Fig. 6a), which was interesting as the S-OPA1 isoform has been shown to protect cells from death by inhibiting mitochondrial release of cytochrome c^37,38^. Another mitochondrial protein, Mitofusin1 and the endoplasmic reticulum protein, Calreticulin were both present at similar levels in naked mole-rat cells compared to human cells (Extended Data Fig. 6 b, c). We next tested if naked mole-rat OPA1 is necessary to confer resilience to oxygen deprivation. Using a shRNA-lentiviral based approach we generated naked mole-rat fibroblasts, in which the OPA1 protein abundance was brought down to levels observed in human and Cape mole-rat cells (Fig. 5e). Crucially when we exposed these cells to extreme hypoxia and compared them to control transfected cells, OPA1 knockdown cells now showed susceptibility to hypoxia equivalent to that seen in human and Cape mole-rat cells (mean T_50_ < 20 h), control transfected cells survived significantly longer (T_50_ ∼ 30 h) (Fig. 5 e, f). Thus, unique genomic rearrangements leading to meaningful structural changes in OPA1 oligomers appear to have evolved at least one molecular mechanism that protects naked mole-rat cells from the consequences of extreme hypoxia.

**Figure 5.**
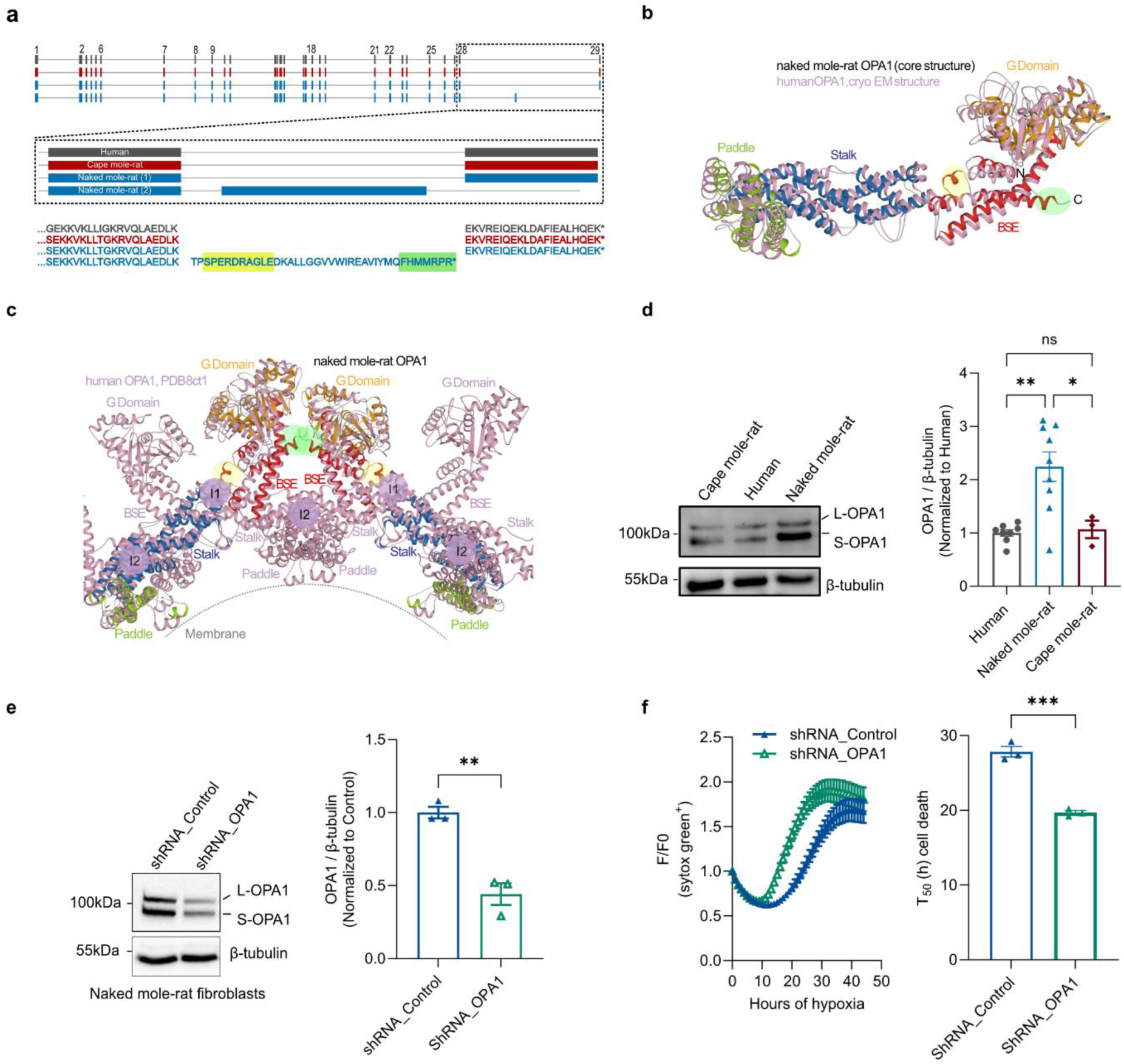
OPA1 is essential for naked mole-rat hypoxia resistance. **(a)** Alignment of transcriptomic sequence of human, naked mole-rat and cape mole-rat OPA1 C-terminus. In yellow and in green is indicated the sequence generating the insert and the C-terminal extension shown in Fig5 b, c. **(b)** Superposition of the naked mole-rat OPA1 core structure and the experimentally determined human OPA1 structure in cartoon presentation. The domains of the naked mole-rat OPA1 core structure are colored orange for the G domain, red for the BSE, blue for the stalk and green for the paddle. The cryo-EM structure of human OPA1 is shown in light pink. The bulky insert in naked mole-rat OPA1 (yellow sphere) and the C-terminal extension (green sphere) are highlighted. **(c)** Cartoon illustration of cryo-EM structure of a membrane bound OPA1 dimers (light pink) and superposition of two naked mole-rat OPA1 molecules reveal new interactions sites formed by the bulky insert (yellow sphere) and the C-terminal extension (green sphere). **(d)** Representative western blot (left) and quantification of OPA1 levels in human, naked mole-rat and cape mole-rat fibroblasts. Total -OPA1 was quantified and normalized to β-tubulin. Data are normalized to human. n is number of experiments, n>3. One-way ANOVA. **(e)** Representative Western blot (left) and quantification (right) of OPA1 and β-tubulin in naked mole-rat cells transduced with control and OPA1 shRNA lentivirus. Data are normalized to control. n is the number of experiments, n=3. **(f)** Mean survival curve (left) and cell death time 50 (T50) (right) in naked mole-rat primary fibroblasts transduced with either control shRNA or OPA1shRNA. Cells are exposed to up to 48 h of hypoxia (1%O2). n is number experiments, n=3. Student t-test. *p < 0.05, **p < 0.01, ***p < 0.001. Data are presented as mean values ± s.e.m.

## Discussion

Here we show that the in vivo anoxia susceptibility of different African mole-rat species scales with sociality. Most importantly, solitary Cape mole-rats were just as susceptible as mice and rats to anoxic challenge. We also show that at the cellular level, naked mole-rat fibroblasts show a remarkable resilience to hypoxia, not seen in fibroblasts taken from the solitary Cape mole-rat, nor in human or mouse fibroblasts. Our mechanistic studies showed that naked mole-rat cells survive hypoxia partly by bypassing mitochondrial oxidative respiration and producing ATP primarily via glycolysis. Naked mole-rat fibroblasts are also capable of withstanding hypoxia by ATP production powered by fructose, a metabolic trick not found in human cells. Naked mole-rat cells are equipped with unusually small mitochondria with sparse cristae and proteomic analysis indicated that naked mole-rat mitochondria are pre-adapted to hypoxia. More importantly, hypoxia initiates mitochondrial fragmentation ending in cell death, but we show here that naked mole-rat mitochondria are highly resistant to fragmentation. Experiments where naked mole-rat cells were forced to utilize mitochondria for energy production (glucose replaced with galactose), led to mitochondrial fragmentation and rapid cell death under hypoxia, thus pinpointing the central role of this organelle in cell survival upon hypoxic stress. We further identified OPA1 as one critical protein, present at high levels in naked mole-rat mitochondria, that has undergone specific structural changes that may have enhanced its protective function. Indeed, we show that reducing naked mole-rat OPA1 levels is sufficient to make naked mole-rat cells similarly susceptible to hypoxia as human or Cape mole-rat cells. These data are consistent with the idea that higher levels of a structurally more robust OPA1 oligomer in naked mole-rat cells can more efficiently hinder the cytochrome c release that initiates apoptosis.

Several studies, including our own ^4^, have indicated that there is altered mitochondrial function in naked mole-rat cells characterized by reduced oxidative respiration^9–11^. However, the molecular mechanisms whereby naked mole-rat cells and tissues resist hypoxic stress have remained elusive. Here we show that specific mitochondrial adaptations, morphological as well as molecular, are central to cellular hypoxia resistance in this species. The fission and fragmentation of mitochondria is a critical step that leads to cell death and we identify one protein, OPA1, as playing a critical role in protecting naked mole-rat cells from hypoxia. OPA1 plays an important role in mitochondrial dynamics and cristae morphogenesis^29^. Deletion of the *Opa1* gene in mice is embryonic lethal^39,40^ and high levels of the protein can be toxic to cells^41,42^. However, moderate over-expression of OPA1 has been shown to be protective e.g. against ischemic damage. In this case, OPA1 is thought to be protective by delaying the release of cytochrome c in response to pro-apoptotic signals^31,38,43^. The long form of OPA1 (L-OPA1) anchored to the inner mitochondrial membrane orchestrates mitochondrial fusion, and its cleavage by the metalloprotease OMA1 and the i-AAA protease Yme1L is induced by stress (e.g. oxidative stress). The accumulation of the short form of OPA1 (S-OPA1) leads to mitochondrial fragmentation, implicating S-OPA1 in fission and mitochondrial quality control essential for cell protection during under pro-apoptotic conditions^41,44,45^. Our finding that both L- and S-OPA1 are higher in small naked mole-rat mitochondria are consistent with these findings. It appears that the eusocial naked mole-rat adapted to regular hypoxic episodes e.g. during sleep by upregulating OPA1, particularly the S-OPA1 which is stress-responsive. However, we find here that it may not just be the levels of OPA1 that are important. We discovered a novel C-terminal exon specifically in the *Opa1* locus which alters the C-terminal peptide sequences of naked mole-rat OPA1. Structural modelling revealed that this novel isoform could form more stable oligomeric forms, compared to the OPA1 in humans, mice or Cape mole-rats. Thus, changes in both OPA1 levels and its filament composition may be protective by hindering cytochrome c release from damaged mitochondria following hypoxia. Although OPA1 clearly plays a key role in protecting from hypoxic stress our data on the mitochondrial proteome will be a rich resource to identify other protective molecular adaptations that enable the extraordinary resistance of naked mole-rat cells to hypoxic stress. We show that by gaining a molecular understanding of the role of mitochondria in protecting from hypoxic stress it may in the future be possible to reengineer human mitochondria for the treatment of hypoxic damage in stroke and heart failure.

## Methods

### Animals

All animal protocols were approved by the University of Illinois at Chicago Institutional Animal Care and Use Committee, the local governmental authorities in Berlin (Landesamt für Gesundheit und Soziales, Berlin), or the Animal Use and Care Committee of the University of Pretoria (EC081-12 & NAS209-2021), Republic of South Africa. Naked mole-rats (*Heterocephalus glaber*) used in this study were kept at the Max-Delbrück Center for Molecular Medicine in Berlin. Naked mole-rats were maintained in a humidity (50-70%) and temperature (30-32 °C) controlled environment, under low illumination levels. A diet of vegetables (primarily sweet potatoes, celery root, carrots and cucumber) was provided daily (*ad libitum*). Animals were housed by colony in a series of custom designed interconnected plastic chambers (Fräntzel Kunstsstoffe, Rangsdorf, Germany). All other Bathyergidae species were housed at the University of Pretoria where animals were housed at room temperature (24-26°C) and humidity (40-60%) in several plastic chambers (1 m× 0.5 m× 0.5 m), with wood shavings and paper toweling provided as nesting material. They were fed a variety of chopped vegetables (primarily sweet potatoes, apples and carrots). C57BL/6N mice were housed with food, water and enrichment available *ad libitum*.

### In vivo experiments

Animals were placed into a clear plastic chamber pre-filled with 100% nitrogen. Thereafter the chamber was infused continuously at 10 liters per minute. Using an Ocean Optics Foxy-PI200 probe, and an Ocean Optics sensor connected to a computer, we measured the fill time, which was, on average, 59.7 ± 2.3 seconds (standard error). Based on the data, we pre-filled the chamber for 120 seconds prior to introducing the animal. Breaths were recorded visually and counted by an observer with a manual counter. “Time Breathing” was determined as the last breath before a 60-second period of no respiration attempts. At that point, animals were removed from the chamber and placed into room air. Each experiment was video recorded to confirm the timing data that was collected in real-time.

### Mitochondria respiration in isolated mitochondria

#### Mitochondria isolation

Adult naked mole-rats and mice (C57BL/6N) had no access to food for at least 2 hours prior to the experiment. Animals were sacrificed by cervical dislocation and the liver was immediately removed, minced with pre-cooled scissors and homogenized using Glass/Teflon Potter Elvehjem homogenizers in isolation buffer (70 mM sucrose, 210 mM mannitol, 5 mM HEPES, 1 mM EGTA, 0.5% BSA, pH 7.2). The mitochondria suspension was transferred into 15 ml Falcon tubes and centrifuged at 1000g for 10 mins. The resulting supernatant containing the mitochondria was collected and centrifuged at 10,000g for 10 mins at 4°C. The mitochondria-enriched fraction was washed with isolation buffer and centrifuged at 10,000g for 5 minutes at 4°C. The final pellet containing mitochondria was suspended in 200-300 μl of isolation buffer and stored on ice. Protein content in mitochondria samples was determined by Bradford protein assay (Biorad 5000001). Freshly isolated mitochondria were used immediately for mitochondrial respiration.

#### Mitochondria respiration measurement

Measurements of mitochondria respiration were monitored with a Hansatech Oxytherm Clark-type O2 electrode connected to a Hansatech chart recorder. The mitochondria suspension was loaded in a sealed chamber filled with respiration buffer (70 mM sucrose, 220 mM mannitol, 2 mM HEPES, 1 mM EGTA, 10 mM KH2PO4, 5 mM MgCl2, 0.2% BSA, pH 7.2), exposed to the surface of a Clark oxygen electrode. Fractional concentrations of oxygen were recorded at 2s intervals at 30°C, which was chosen because it was nearest to physiological conditions for both naked mole-rats and thermoneutral mice. The O_2_ electrode was calibrated using air-saturated water and sodium dithionite according to the manufacturer’s protocol. Freshly isolated mouse mitochondria in a concentration 0,5 mg/ml were incubated in respiration buffer. The following substrates were used to follow mitochondrial respiration: 10 mM malate and 10 mM glutamate (complex I substrates), or 10 mM succinate (complex II substrate) with 1µM rotenone (complex I inhibitor, Sigma-Aldrich R8875). After measuring oxygen consumption (State 3) upon addition of 200 nmol ADP (Sigma Aldrich, 58-64-0), 1 μM oligomycin A (Th.Geyer, 75351) was added to block mitochondrial ATP synthase to examine the residual respiration reflecting proton leak (state 4). Maximum oxygen consumption was also measured in the presence of 1 μM carbonylcyanide-p-trifluoromethoxyphenylhydrazon (FCCP, Sigma Aldrich C2920).

The respiration rate was calculated using Hansatech Oxytherm software. The average of the 30 values before the following addition was calculated and the rates of O2 consumption were expressed in nmol O2/mg mitochondrial protein/minute of respiration. The respiratory control ratio (RCR) is the ratio between State3 (ADP-induced oxygen consumption) and State4 (oligomycin-induced oxygen consumption). The respiratory capacity ration is the ratio between the oxygen consumption rate upon oligomycin and the rate upon FCCP addition.

### Naked mole-rat primary fibroblasts isolation

Naked mole-rat and Cape mole-rat fibroblasts were isolated as described^46^. Briefly, skin and kidneys from neonatal, 30 and 60-day-old naked mole-rat were collected after animal decapitation. Kidneys from neonatal Cape mole-rat pups were collected at the University of Pretoria. The tissues were immediately placed in the Cell Isolation Medium (DMEM high glucose, Gibco #41966029) supplemented with 200 units/ml Penicillin, 200µg/ml Streptomycin (Gibco #15140122) and 200µg/ml Primocin (InvivoGen # ant-pm-2). Skin came from either the underarm area and was cleared of any fat or muscle tissue and sprayed with 70% ethanol before collection. Adrenal glands were removed from the kidneys. All tissues were then washed twice with cold PBS and finally minced with sterile scalpels. Cape mole-rat tissues were then placed in BamBanker medium (Nippon Genetics #BB03-NP) and freeze at −80 °C. The samples were then shipped to Max Delbrück Center for Molecular Medicine in Berlin where the cells were isolated and cultured. Minced tissues, freshly obtained from naked mole-rats or cryopreserved in BamBanker medium, were transferred in 5ml of NMR Cell Isolation Medium containing 500µl of Cell Dissociation Enzyme Mix: 10mg/ml Collagenase (Roche #11088793001) and 1000Units/ml Hyaluronidase Sigma #H3506) in DMEM high glucose (Gibco #41966029) and incubated at 37°C for 2 hours for the skin and 45 minutes for kidneys. Each tissue was briefly vortexed every 30 minutes to aid cell dissociation. Cells were then pelleted by centrifuging at 700 g for 5 minutes and resuspended in NMR Cell Culture Medium (DMEM high glucose (Gibco #41966029) supplemented with 15% fetal bovine serum (Gibco), 1x non-essential amino acids (Gibco # 11140050), 100units/ml Penicillin, 100µg/ml Streptomycin (Gibco #15140122) and 100µg/ml Primocin (InvivoGen # ant-pm-2). This cell suspension was passed through 70µm filter (Corning #352350) and seeded on cell culture dishes. Naked mole-rat fibroblasts were placed in a humidified 32°C incubator with 5% CO2 and 5% O2; Cape mole-rat fibroblasts were placed in an humidified 37°C incubator with 5% CO2. Medium was changed the day after and then every 2-3 days until the cells got confluent.

### Naked mole-rat, Cape mole-rat, mouse and human primary fibroblasts culture

Human (neonatal, Innoprot #P10856), mouse (embryonic, Gibco #A34180), naked mole-rat and Cape mole-rat primary fibroblasts were cultured in glucose or galactose medium as indicated. Medium was changed every 2-3 days and cells split once they reached 80-90% confluency. Human, mouse and cape mole-rat fibroblasts were kept in a humidified 37°C incubator with 5% CO2. Naked mole-rat fibroblasts were kept in a humidified 32°C incubator with 5% CO2, 5% O2.

Naked mole-rat kidney primary fibroblasts were used for most of the experiments, where indicated skin primary fibroblasts were used.

#### Normoxia conditions

Human, mouse and cape mole-rat fibroblasts: 37°C, 5% CO2. Naked mole-rat fibroblasts: 32°C, 5% CO2, 5% O2.

#### Hypoxia conditions

Human, mouse and cape mole-rat fibroblasts: 37°C, 5% CO2, 1% O_2_ Naked mole-rat fibroblasts: 32°C, 5% CO2, 1% O2.

#### Glucose medium

DMEM high glucose, pyruvate (Gibco # 41966029) supplemented with 15% fetal bovine serum (Gibco), 1x non-essential amino acids (Gibco # 11140050), 100units/ml Penicillin, 100µg/ml Streptomycin (Gibco # 15140122) and 100µg/ml Primocin (InvivoGen # ant-pm-2).

#### Galactose medium

DMEM no glucose (Gibco # 11966025) supplemented with 1% fetal bovine serum (Gibco), 10mM D-(+)-Galactose (Sigma-Aldrich), non-essential amino acids (Gibco # 11140050), 1mM sodium pyruvate (Gibco #11360039), 100units/ml Penicillin, 100µg/ml Streptomycin (Gibco # 15140122) and 100µg/ml Primocin (InvivoGen # ant-pm-2). Cells were culture in galactose medium at least 5 days before starting the experiment.

#### Fructose medium

DMEM no glucose (Gibco # 11966025) supplemented with 1% fetal bovine serum (Gibco), 10mM Fructose (Sigma-Aldrich), non-essential amino acids (Gibco # 11140050), 1mM sodium pyruvate (Gibco #11360039), 100units/ml Penicillin, 100µg/ml Streptomycin (Gibco # 15140122) and 100µg/ml Primocin (InvivoGen # ant-pm-2). Cells were culture in fructose medium at least 5 days before starting the experiment.

### Transmission Electron microscopy

#### Liver

Adult naked mole-rats and mice (C57BL/6N) were anesthetized and perfused with a solution of 4% paraformaldehyde [v/v] and 1.25% glutaraldehyde [v/v] in phosphate buffer. The liver was removed, cut into pieces of 1mm and placed in 2.5% GA in phosphate buffer overnight at 4°C. For cryo sectioning, samples were infiltrated with 2.3 M sucrose. Samples were sectioned at −110°C with 60nm thickness (Ultra cut Leica Microsystems, Germany). To better visualize the inner membrane structures ultrathin cryosections were contrasted and stabilized with a mixture of 3% [w/v] tungstosilicic acid hydrate (Sigma-Aldrich) and 2.5% [w/v] polyvinyl alcohol (Sigma-Aldrich)^47^ . Cryo sections were examined with a FEI Morgagni electron microscope, images were taken with a Morada CCD camera and the iTEM software (Olympus Soft Imaging Solutions GmbH, Münster, Germany). Mitochondrial area and number of cristae/mitochondria area were analyzed with Fiji.

#### Human and naked mole-rat primary fibroblasts

Cells were seeded on carbon-coated sapphire discs and after 24 hours cells are kept in normoxia (normal culture conditions) or exposed to 4 hours of hypoxia (<1%O_2_).

Immediately after cultivation or hypoxia treatment the primary fibroblasts were cryo-fixed with 20 % [w/v] ficoll in cell culture media using the EM ICE system (Leica Microsystems, Germany). For freeze substitution, samples were incubated in a solution containing 1% H₂O [v/v], 1% glutaraldehyde [v/v] (Electron Microscopy Sciences, USA), 2% osmium tetroxide [w/v] (Roth, USA) in anhydrous acetone (Sigma-Aldrich, USA)) at −90°C. The Automatic Freeze Substitution system (AFS 2, Leica Microsystems, Germany) was used with the following program: −90°C for 36 hours, −90°C to −50°C over 8 hours, −50°C to −20°C over 6 hours, −20°C for 12 hours, and −20°C to 20°C over 3 hours. Discs were further processed at room temperature by incubation in 0.1% [w/v] uranyl acetate (UA) in acetone. Infiltration with Durcupan in acetone was conducted with increasing concentrations of 30%, 70%, 100% Durcupan each for 1 hour at room temperature. The final Durcupan mixture comprised of 20g component A, 20g component B, 0.4g component C, and 0.48g component D (Sigma-Aldrich, USA). For the minimal resin embedding of cell monolayers on sapphire discs we used a modified approach^48^. To achieve a thin resin layer the incubation in the final step of 100% durcupan was done at 50°C. The discs were cleaned of excess resin, transferred to a microscopy slide, and incubated in acetone vapors for 10 minutes at room temperature and 15 minutes at 50°C, followed by a centrifugation for 2 minutes at 1000g. Discs were placed with a tilted orientation in an incubator and polymerized for 2-3 days at 60°C.

### FIB-SEM

For 3D analysis, minimal resin embedded cells were mounted onto the SEM stabs and sputter coated with 50 nm carbon (Carbon Coater). 3D stacks of cells were acquired using Helios 5CX FIB-SEM and the Auto Slice & View platform [Thermo Fisher Scientific, USA]. SEM imaging conditions −0,34nA, 2kV, 3,37x4,6x10 nm voxel. Image processing and mitochondria segmentation was performed using Microscopy Image Browser v.28454 and its 2,5D Deep learning procedure. Mitochondria network was manually proved to make sure that all mitochondrial objects are correctly separated from each other. 3D visualization was done using Imaris^49,50^.

### Cell viability

Human, mouse, Cape mole-rat, naked mole-rat, naked mole-rat shRNA-control and naked mole-rat shRNA OPA1 fibroblasts (1.5×10^5^ cells/well) were seeded on black 96 well plate. After 24 hours cells were loaded with 200nM SYTOX™ Green Nucleic Acid Stain (Invitrogen #S7020) and placed in Cytation C10 (Agilent) at 37 °C (human, mouse and Cape mole-rat) or 32°C (naked mole-rat). Here GFP signal and brightfield images were simultaneously acquired every hour for 24 hours (for human, mouse and Cape mole-rat) or 48 hours (for naked mole-rat). After one baseline acquisition, the O_2_ level was set to 1% O_2_ using the gas controller (Agilent). F/F0 was calculated, where F0 is the mean of the first two fluorescence measurements. F% was used to calculate the T_50_(h).

### Mitochondrial morphology analysis

#### Fixed cells

Human and naked mole-rat fibroblasts are seed in 15mm diameter glass coverslip (30×10^5^ cells/coverslip). After 24 hours cells were kept in normoxia (standard culture conditions) or placed in a hypoxia chamber (Stem Cell # #27310) for 2, 4, 8 and 24 hours of hypoxia (<1%O_2_). Cells kept in normoxia or exposed to hypoxia were fixed in 4% PFA (15 min) and then washed 3 times with PBS. Cells were then permeabilized with 0.25% Triton X-100 in PBS (10 min) and blocked with a PBS solution containing 2% BSA for 1 hour. Cells were incubated overnight at 4°C with primary antibody diluted in blocking solution (dilution 1:100 Tom20 F-10 Santa Cruz #sc-17764). The following day, cells were washed 3 times with blocking solution and incubated for 45 min at RT with secondary antibody (1:300 dilution in blocking solution; Invitrogen #A-11029). Coverslips were washed 3 times (5 min) with the blocking solution and with PBS (10 min). Cells were then incubated with dapi for 10 minutes. They were finally mounted using Dako (#S3023). Images were collected with a Zeiss LSM700 confocal microscope.

#### Live cells

Human, naked mole-rat and Cape mole-rat fibroblasts were transduced with lentivirus pLYS1-FLAG-MitoGFP-HA (Addgene plasmid #50057; was a gift from Vamsi Mootha^51^) which contains the pore-forming subunit of the mitochondrial calcium uniporter to target the label to the mitochondria. After 72 hours the medium was changed to fresh culture medium and cells were split once they reached 80-90% confluency. For imaging, cells were seeded in µ-Slide 8 Well (Ibidi). After 24 hours, cells were placed in Cytation C10 (Agilent) at 37 °C (human and Cape mole-rat) or 32°C (naked mole-rat). Here images were acquired (60x objective – confocal) before (normoxia) and after 4 hours of hypoxia (<1%O_2_). The levels of oxygen were controlled by the gas controller (Agilent).

Mitochondrial morphology analysis was performed using the Fiji plugin Mitochondrial Analyzer as previously described^52^. The cell area, mitochondrial number, area and perimeter were considered.

### Total cellular ATP measurements

Human and naked mole-rat fibroblasts (1.5×10^5^ cells/well) are seeded in a white 96-well plate in glucose or galactose medium (see above); after 24 hours total ATP content in each well was measured. For hypoxia, cells were placed in cytation5 at 37 °C (human) or 32°C (naked mole-rat) and the O_2_ level was set to 1%O_2_ using the gas controller (Agilent). After 4 hours of hypoxia, total cellular ATP levels were measured. ATP levels were measured with the luciferin/luciferase assay ATPlite 1 step (PerkinElmer, 6016736), and luminescence was measured at cytation5 (Agilent). Luminescence was normalized to µg of proteins.

### Mitochondrial Transmembrane Potential Measurements

Mitochondrial membrane potential (ΔΨ) was detected by tetramethyl rhodamine methyl ester (TMRM) fluorescent dye.

Human and naked mole-rat fibroblasts (1.5×10^5^ cells/well) were seeded µ-Slide 8 Well (Ibidi). After 24 hours, cells were loaded with 10 nM tetramethyl rhodamine methyl ester (TMRM) supplemented with Cyclosporin H (2 mg/ml; Sigma Aldrich, SML1575) to inhibit multidrug-resistance pumps, which could affect TMRM loading. Cells were placed in Cytation C10 at 37 °C (human) or 32°C (naked mole-rat) and cells were imaged using 40X objective with mCherry filter. Images were collected every 120s (300 ms exposure) for 40 minutes. Where indicated, 10 mM FCCP was added to assess the correct distribution of the dye. Images were analyzed with ImageJ.

### Western Blot

#### Preparation of protein extracts

Human, naked mole-rat and Cape mole-rat fibroblasts were placed in ice and washed once with cold PBS. A cell scraper was used to collect cells in RIPA buffer (50 mM Tris, 150 mM NaCl, 1% Triton X-100, 0.5% deoxycholic acid, 0.1% SDS, protease and phosphatase inhibitor cocktails (Roche), pH 7.5). Homogenates were incubated on ice for 30 min, centrifuged at 13000g for 15 min at 4°C and the supernatant was collected. Protein concentration was measured by a BCA protein assay kit (Thermo scientific #23227). 10-30 mg of proteins were loaded onto polyacrylamide gels (8-12%) and immunoblotted as previously described ^53^.

Membranes were incubated with primary antibodies in 5% milk (α-Mitofusin1, rabbit, Proteintech, #13798-1-AP, 1:500; α-Calreticulin, Thermo Fisher, #PA 3-900, 1:1000; α-β-tubulin, mouse, Sigma Aldrich #T4026; α-β-Actin, mouse, Sigma Aldrich #A1978; α-OPA1, mouse, BD Transduction #612607, 1:1000; α-Vinculin, rabbit, Abcam #ab73412, 1:500) overnight at 4°C. After overnight primary antibody incubation, secondary species-specific HRP-coupled antibodies have been used. The proteins were visualized by the chemiluminescent reagent ECL (Life Technologies, # 32106) at the ChemiDoc MP (Biorad).

### ATP Synthase

Isolated liver crude mitochondria from adult naked mole-rat or mouse (C57BL/6N) were resuspended in 50 mM NaCl, 50 mM imidazole/HCl, pH 7.0, 2 mM 6-aminocaproic acid, 1 mM EDTA at a final concentration of 10 μg/μl, in presence of the indicated amount of digitonin and subjected to an ultraspin at 100,000 x g for 30 min at 4°C. The resulting supernatant was collected, supplemented with Blue Coomassie G-250 (5% G-250 Sample Additive, Invitrogen) and loaded onto a Native-PAGE 3–12% gel. The native gel was then stained with Coomasie Blue or transferred to a PVDF membrane and subjected to western blot for subunit c of ATP synthase (ab181243, Abcam).

### PCR and agarose gel

Total RNA from naked mole-rat and human fibroblasts and from naked mole-rat tissues was extracted using the ReliaPrep™ RNA Miniprep System (Promega Corporation, #Z6010 and #Z6110), followed by retrotranscription to cDNA using the GoScript™ Reverse Transcriptase kit (Promega Corporation, # A5003) according to the manufacturer’s instructions. DNA amplification was performed using Q5® Hot Start High-Fidelity DNA Polymerase (New England Biolabs, #M0493) following the manufacturer’s protocol. Thermal cycling conditions were optimized as follows: initial denaturation at 98°C for 30 s, followed by 35 cycles of 98°C for 10 ss, annealing at a gradient of 65-68°C for 20 s, and extension at 72°C for 45 s, with a final extension at 72°C for 2 minutes. PCR products were then separated on a 1.5% agarose gel. The PCR products were purified using the Wizard® SV Gel and PCR Clean-Up System (Promega Corporation, #A9281). To prepare the blunt-ended PCR products for cloning, an A-tailing reaction was performed using Taq DNA Polymerase, recombinant (Invitrogen, # 10342020) and 100 mM dATPs. The A-tailed PCR products were then cloned into the pGEM®-T Easy Vector (Promega Corporation, # A1360) according to the manufacturer’s instructions. The ligated vectors were transformed into NEB 5-alpha competent E. coli (New England Biolabs, Catalog Number C2987) using the IPTG/X-gal system for blue/white screening. Transformed cells were plated on LB agar plates containing IPTG and X-gal, and white colonies were selected for further analysis. Plasmid DNA was extracted using the PureYield™ Plasmid Miniprep System (Promega Corporation, # A1223) according to the manufacturer’s protocol. The purified plasmids were then sent for Sanger sequencing.

Primers:

F1_OPA1 5’-GTTACAGACTTGGTCAGTCAAATGG -3’

R1_OPA1 5’-CTTGTAGGGTCTCCCAAGCAAC-3’

F2_OPA1 5’-CACAGTCCGGAAGAACCTTGAATC-3’

R2_OPA1 5’-CTACCGAGGTCTCATCATATGGAA-3’

### Lentivirus production

HEK293T cells were cultured in DMEM high glucose, pyruvate (Gibco #41966029) supplemented with 10% fetal bovine serum (Gibco), 1x non-essential amino acids (Gibco #11140050), 100units/ml Penicillin, 100µg/ml Streptomycin (Gibco #15140122) were transfected using a calcium phosphate–based transfection method with 15μg of shuttle vector and 10.5 or 4.5μg of the helper plasmids psPAX2 and pMD2.G, respectively. Twelve hours after transfection, the culture medium was replaced with a fresh medium. Supernatants containing viral particles were collected 48 and 72 hours after transfection, filtered to remove cell debris, and concentrated to 30-fold using Amicon tubes (Ultra-15, Ultracel-100k; Merck Millipore). Aliquots were flash-frozen in liquid nitrogen and stored at −80°C until use.

### Label-free proteomics on liver

#### Preparation of proteins and peptides for mass spectrometry

Adult naked mole-rats and mice (C57BL/6N) had no access to food for at least 2 hours prior to the experiment. Animals were killed by decapitation, and tissues were isolated and immediately frozen in liquid nitrogen. For protein extraction, samples were resuspended in urea buffer (8 M Urea, 100 mM Tris-HCl, pH 8.25) containing 100 µl of zirconium beads and homogenized on a Precellys 24 device (Bertin Technologies) using two cycles of 10 sec at 6000 rpm. After a centrifugation step to remove beads and tissue debris, protein concentration was measured by Bradford colorimetric assay and 100 µg of protein from each sample were taken for enzymatic digestion. Briefly, the disulfide bridges of proteins were reduced in 2mM DTT for 30 minutes at 25 °C and the resulting free cysteines alkylated in 11 mM iodoacetamide for 20 minutes at 25 °C in the dark. Samples were then incubated with 5 μg of LysC (Wako) and incubated for 18 hours with gentle shaking at 30 °C. After LysC digestion, the samples were diluted 3 times (v/v) with 50 mM ammonium bicarbonate solution, 7 μl of immobilized trypsin (Applied Biosystems) were added and samples were incubated 4 hours under rotation at 30 °C. 18 μg of the resulting peptide mixtures were desalted on StageTips as described previously^54^ and reconstituted to 20 μl of 0.5 % acetic acid in water.

#### LC-MS/MS analysis

Five microliters were injected in duplicate on a UPLC system (Eksigent), using a 240 minutes gradient ranging from 5% to 45% of solvent B (80% acetonitrile, 0.1 % formic acid; solvent A= 5 % acetonitrile, 0.1 % formic acid). For the chromatographic separation 30 cm long capillary (75 µm inner diameter) was packed with 1.8 µm C18 beads (Reprosil-AQ, Dr. Maisch). On one end of the capillary nanospray tip was generated using a laser puller, allowing fretless packing. The nanospray source was operated with a spay voltage of 2.1 kV and an ion transfer tube temperature of 220 °C. Data were acquired in data-dependent mode, with one survey MS scan in the Orbitrap mass analyzer (60000 resolution at 400 m\z) followed by up to 20 MS\MS scans in the ion trap on the most intense ions. Once selected for fragmentation, ions were excluded from further selection for 40 seconds, in order to increase new sequencing events.

#### MaxQuant data analysis

Raw data were analyzed using the MaxQuant proteomics pipeline v2.4.2.0 and the built in the Andromeda search engine^55–56^ with the Uniprot Mouse proteome database (updated on 5^th^ January 2023) Carbamidomethylation of cysteines was chosen as fixed modification, oxidation of methionine and acetylation of N-terminus were chosen as variable modifications. Two missed cleavage sites were allowed and peptide tolerance was set to 7 ppm. The search engine peptide assignments were filtered at 1% FDR at both the peptide and protein level. The ‘second peptide’ feature and ‘match between runs’ feature (time window= 0.7min) were enabled, while other parameters were left as default. Fold-change was calculated as LFQ intensity naked mole-rat/LFQ intensity mouse <0.5 and >2 was considered for GO-Term analysis^57^. GO-Term analysis was performed in naked mole-rat up and downregulated proteins using ShinyGO^58^.

### TMT-labeling proteomic on crude mitochondria from fibroblasts

#### Crude mitochondria isolation

Human and naked mole-rat fibroblasts were kept in normoxia or exposed to 4 hours of hypoxia (1%O_2_). Cells were then washed 2 times with cold PBS and collected with a cell scraper. Cells were collected in a falcon tube and centrifuged for 5min, 800g, 4°C. The pellet was resuspended in cold PBS and cells were centrifuged for 5 min, 800g, 4°C. The pellet was resuspended in Buffer M (225mM Mannitol, 75mM Sucrose, 10mM Hepes, 2mM EGTA) and transferred in a glass tube. Here cells were homogenized with a Teflon Potter Elvehjem homogenizer (900rpm, 30 strokes) in ice. The suspension was centrifuged for 5 min, 800g, 4°C. The pellet was discarded, and the supernatant was collected and centrifuged 10 min, 8000g, 4°C. The pellet containing the crude mitochondrial fraction was used.

#### Proteolytic digestion

Crude mitochondria resuspended in TH buffer (10 mM KCl, 10 mM Hepes-KOH pH 7.4, 250 mM Threalose) were supplemented with 8 M Urea and lyzed using a Bioruptor (10 min, 30 s on, 30 s off). All subsequent steps were performed independently for all samples at 37 °C. Proteins were reduced with 5 mM DTT for 30 min, alkylated with 40 mM CAA for 1 h and proteolytically digested with 1:200 enzyme:substrate (wt/wt) LysC for 2 h. After dilution of the sample to 2 M Urea, a tryptic digest with 1:100 enzyme:substrate (wt/wt) Trypsin was performed overnight. The digestion was stopped with 1% FA. Peptides were desalted using C8 SepPak (Waters) and dried in a SpeedVac.

#### TMT labeling

Dried peptides were resuspended in 50 mM TEAB pH 8.5. 320 µg TMT 10-plex reagent (Thermo Fisher Scientific) was added to 200 µg peptide and incubated for 1 h at room temperature. The reaction was quenched by the addition of 66 mM Tris-HCl pH 8 for 15 min at room temperature. All samples were first combined before excess ACN was removed by vacuum centrifugation. TMT-labeled peptides were desalted using C8 SepPak and dried in a SpeedVac.

#### Strong cation exchange chromatography

500 µg of the TMT-labeled peptides were resuspended in 0.05% FA, 20% ACN and loaded onto a PolySULFOETHYL A column (100 mm x 2.1 mm, 3 µm particle size, PolyLC INC.) with an Agilent 1260 Infinity II system and separated using a 90 min gradient with increasing NaCl concentration. Selected fractions were resuspended with 0.1% FA and desalted using C8 StageTips.

#### LC-MS/MS analysis

Desalted fractions were resuspended in 1% ACN, 0.05% TFA. Reverse-phase separation was performed on a Dionex UltiMate 3000 system (Thermo Scientific) connected to a PepMap C18 trap-column (0.075 mm x 50 mm, 3 μm particle size, 100 Å pore size, Thermo Scientific) and an in-house packed C18 column (Poroshell 120 EC-C18, 2.7 μm particle size, Agilent Technologies) at 250 nL/min flow with increasing ACN concentration. All fractions were analyzed on an Orbitrap Fusion Lumos mass spectrometer with FAIMS Pro device (Thermo Scientific) and Instrument Control Software version 4.0. Fractions were first analyzed with an extensive quantitative cross-linking mass spectrometry method. Then, fractions were further combined and measured with a standard quantitative proteomics acquisition strategy.

|  |  | method (1) | method (2) |
| --- | --- | --- | --- |
| Global | Gradient length | 180 min | 120 min |
|  | Cycle time (mode) | 2 s (top speed) | 2 s (top speed) |
|  | FAIMS compensation voltages | -50, -60, -75 | -50, -65, -85 |
| MS1 | MS1 detector | Orbitrap | Orbitrap |
|  | MS1 resolution | 120,000 | 120,000 |
|  | scan range | 375 – 1,600 | 400 – 1,600 |
|  | maximum injection time | 50 ms | 246 ms |
|  | AGC target | 400,000 | 400,000 |
| Quadrupole | charge filter | 4 – 8 | 2 – 6 |
| | dynamic exclusion | 60 s (± 10 ppm) | 60 s (± 10 ppm) |
| MS2 | MS2 detector | Orbitrap | Orbitrap |
|  | MS2 resolution | 60,000 | 50,000 |
|  | isolation window | 0.7 m/z | 0.7 m/z |
|  | scan range | first mass 110 m/z | first mass 110 m/z |
|  | normalized collision energy | 33%,36%,40% | 38% |
|  | maximum injection time | 118 ms | 86 ms |
|  | AGC target | 100,000 | 125,000 |

#### Database search

RAW files were converted to mzML using ProteoWizard msconvert ^59^. Proteins were identified and quantified using MSFragger v3.9 in fragpipe v20.1^60^. Search parameters were as follows: precursor mass tolerance −20 ppm to 20 ppm, fragment mass tolerance 20 ppm, mass recalibration on, enzyme specificity Trypsin, maximum 2 missed cleavages, peptide length 7 – 50. Variable modifications: TMT6 (+ 229 Da) on serine and peptide N-termini, oxidation of methionine (+ 16 Da), acetylation of protein N-termini (+ 42 Da). Fixed modifications: TMT6 on lysine, carbamidomethylation of cysteine (+ 57 Da). Data were searched against the human proteome retrieved from UniProt on 2021-4-7 and common contaminants. Isobaric quantification was performed using the TMT-Integrator with default settings. Abundances were log2-transformed.

#### Differential expression analysis

The TMT-Integrator output has been further analyzed in R to generate volcano plots. The 0.5% missing values have been imputed using random forest imputation (R package missRanger). Data were median normalized before computing log2-fold changes and p-values (R package limma). Principle component analysis was performed using the R package factoextra. Volcano plots were created using the R package *EnhancedVolcano*. Venn diagrams were created using R packages *VennDiagram* and *eulerr*. GO-Term analysis was performed in up and downregulated proteins (±1.5 fold) using ShinyGO^58^.

### OPA1 alignment

Naked mole-rat transcriptomic data was obtained as raw reads from the National Center for Biotechnology information (NCBI) BioProject PRJNA283581^61^. Raw reads were cleaned using Trimmomatic ^62^ as part of the transcriptome assembly package Trinity^63^, which was used to assemble the transcriptome. In parallel, cleaned reads were mapped using STAR^64^, to the nnotated Ensembl naked mole-rat genome (GCA_944319715.1). Mapped reads and sequence visualization was done in IGV ^65^. Sequences of the isoforms of OPA1 were found using BLAST^66^, using the annotated transcriptomes as reference.

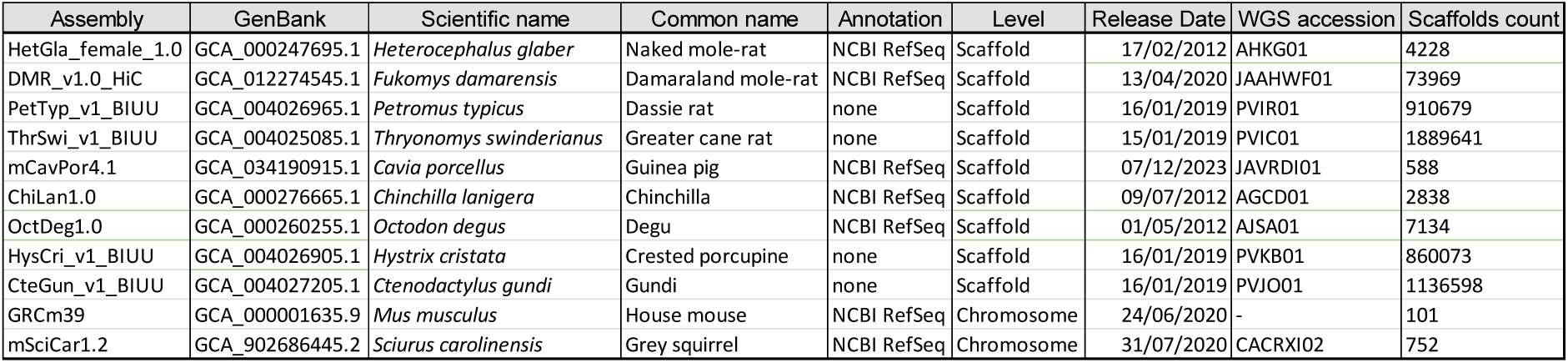

Cape mole-rat and other African mole-rat sequences were obtained from published transcriptomic data ^67,68^. Human and mouse sequences were obtained from NCBI (see table below). Sequence alignments were performed using MAFFT ^69^.

| Species | Accession number | Gene ID | Notes |
| --- | --- | --- | --- |
| *Heterocephalus glaber* | XM_004834760.3 | 101698766 | Human-like isoform |
| *Heterocephalus glaber* | XM_021255509.1 | 101698766 | Naked mole-rat specific isoform |
| *Homo sapiens* | NM_015560.3 | 4976 |  |
| *Homo sapiens* | XM_047448206.1 | 4976 | Isoform with extra exon |
| *Mus musculus* | NM_001199177 | 74143 |  |

NCBI accession numbers of sequences used for comparison.

### Exonic structure investigation

The exonic structure and genomic DNA of the species with RefSeq^70^ annotations were obtained from NCBI (see table below). For the species without annotation, the genome was queried using BLAST using the naked mole-rat sequence as a query. For the species with RefSeq annotation, their respective exons, with the addition of the naked mole-rat specific exon, were mapped to the genomic DNA using the Geneious (https://www.geneious.com) “map to reference” tool, using the highest sensitivity with default parameters (maximum mismatches per read: 50%). The guinea pig exons with the additional naked mole-rat specific exon were used for the species without available annotations. When the map to reference tool did not return any hits, the naked mole-rat specific exon was aligned to the intron using MAFFT. In all cases where the naked mole-rat specific exon was found, it was located in the same region of the same intron as the naked mole-rat.

### ShRNA design

For OPA1 shRNA, the following sequence was designed and used to construct a nuclear localization sequence (NLS)-RFP–containing lentivirus-mediated RNA interference vector targeted to OPA1. shOPA1 5’-GATCCCCACAGTCCGGAAGAACCTTGTTCAAGAGACAAGGTTCTTCCGGACTGTTTTTTGGAAATTAAT -3’.

### OPA1 knockdown

Naked mole-rat fibroblasts were transduced with lentivirus (NLS)-RFP -shRNA-Control or OPA1. After 72 hours the medium was changed to fresh culture medium and cells were split once they reached 80-90% confluency. 20 days after transduction cells were used for hypoxia cell viability experiments or collected for western blot analysis.

### Alpha fold naked mole-rat OPA1 structure prediction

The nmrOPA1 structure was predicted with AF3^34^. The core structures (BSE, stalk, paddle and G domain) were superimposed with Chain G out of the OPA1 assembly (PDB 8ct1, ^35^) using Coot from the ccp4 program suite^71^. Reformatting of the files was done in Chimera^72^. For illustration of the function of the new structural elements in an OPA1 filament, two nmrOPA1 molecules were superimposed in Coot with chains A and I of PDB 8ct1, respectively. Images of the structures were generated with The PyMOL Molecular Graphics System, Version 2.0, Schrödinger, LLC. and labelled in Adobe Illustrator. Software, Colab Alphafold score visualizer, Chimera 1.17, Pymol 2.5.5, Coot 0.8.1, Adobe Illustrator CS6.

## Data availability

Label free proteomic:

The mass spectrometry proteomics data have been deposited to the ProteomeXchange Consortium via the PRIDE^73^ partner repository with the dataset identifier PXD053475“.” Reviewer credentials: Log in to the PRIDE website using the following details:

Project accession: PXD053475

Token: dB8KykwOBdXQ

TMT Proteomic:

1) Data availability: Mass spectrometry RAW files, FragPipe results, experimental annotation and partial analysis outputs were deposited in jPOST^74^ and ProteomeXchange with identifiers JPST003176 and PXD053190. Reviewer credentials:

https://repository.jpostdb.org/preview/103369750966712f80d297b

Access key 8952.

2) Code availability: The R scripts used for processing and visualization of quantitative proteomic data are available at Zenodo (DOI: 10.5281/zenodo.11992058).

## Acknowledgements.

We thank Franziska Bartelt for technical help. The pLYS1-FLAG-MitoGFP-HA was a gift from Vamsi Mootha. We thank Markus Landthaler for the constructive discussion on the analysis of the unique naked mole-rat exon sequence. We thank members of the Lewin lab for constructive comments on the manuscript.

## Author contributions

Conceptualization: A.R. and G.R.L.

Cell culture: A.R with the help of K.P.

Hypoxia cell viability experiments, mitochondrial morphology and functional analysis, molecular biology and lentivirus production: A.R.

Electron microscopy data acquisition and FIB-SEM analysis: S.K, D.P. and B.P.

TEM analysis: A.R., T.K. and V.B.

Liver OCR: T. K., V.B. and A.R.

OPA1 Alignment: D. A M.

OPA1 RT-PCR: K.P.

Fibroblasts TMT proteomics: M.R. and F.L.

Liver proteomic: G.M. and S.K.

ATP synthase BNEG: M.C., L.T and P.B.

African mole-rat species were provided by D.W.H. and N.C.B.

Hypoxia in vivo experiments: D.W.H, N.C.B., T.J.P., J.R., D.O. and G.R.L.

OPA1 structure prediction: K.F. and O.D.

Writing: A.R. and G.R.L., with input from all authors.

Supervision and funding: G.R.L.

## Extended data figures

**Extended data Fig 1.**
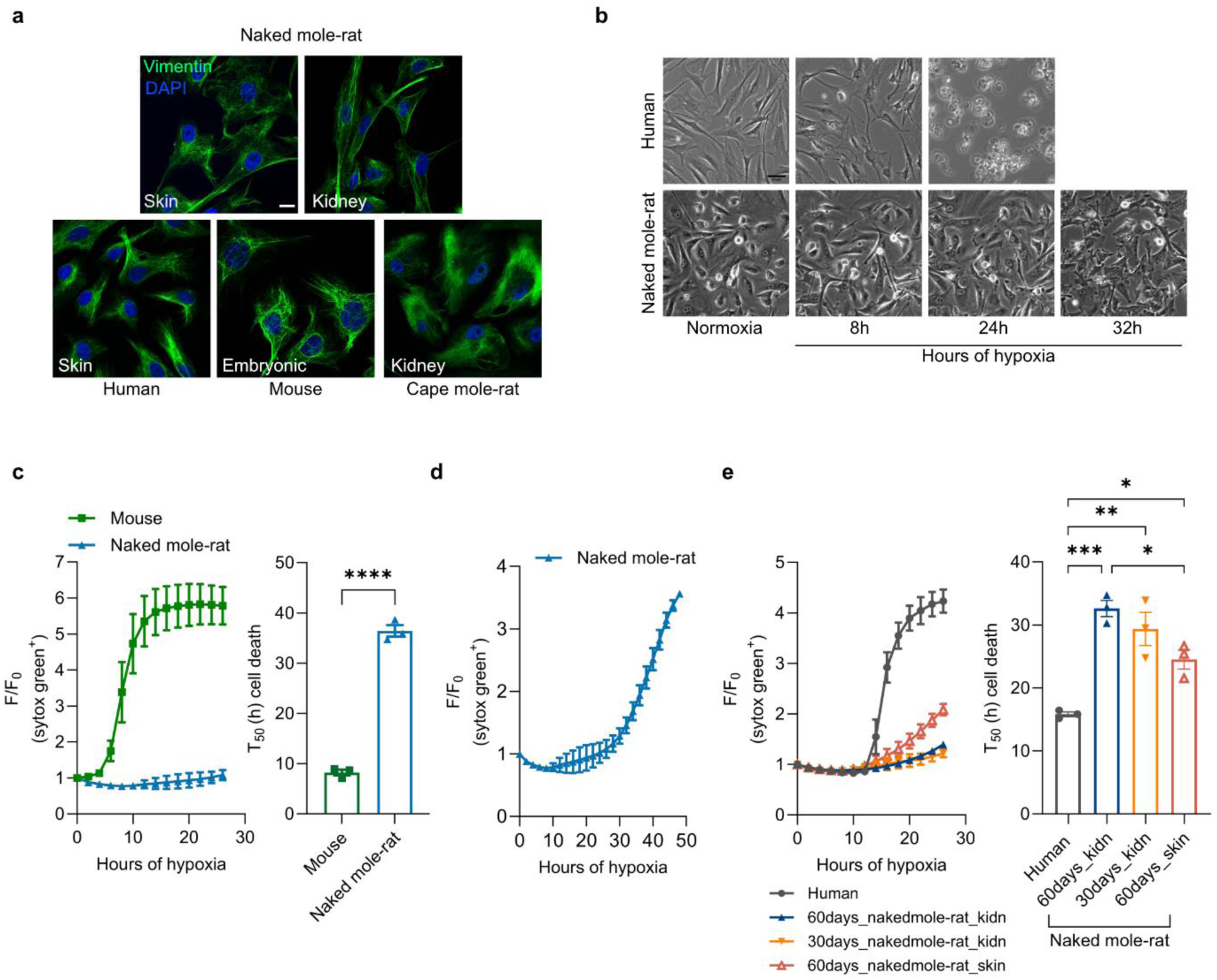
Primary naked mole-rat fibroblasts survive long periods of oxygen deprivation. **(a)** Representative pictures of primary human (skin), mouse (embryonic), naked mole-rat (skin and kidney) and cape mole-rat (kidney) fibroblasts stain with fibroblast specific marker, Vimentin (green) and Dapi (blue). Scale bar 10 µm. **(b)** Representative pictures of human and naked mole-rat cells exposed to hypoxia (1%O_2_). The quantification is shown in Fig 1c and Extended Data Fig. 1c. Scale bar 10 µm. **(c)** Mean survival curve (left) and cell death time 50 (T_50_) in mouse and naked mole-rat primary fibroblasts exposed to 24 h of hypoxia (1%O_2_). Naked mole-rat data (light blue curve and bar) are the same as in Fig 1c. **(d)** Up to 48 h mean survival curve of the viability trace in Fig. 1c of naked mole-rat fibroblasts. Cell death time 50 (T_50_) quantification is shown in Fig. 1c. **(e)** Mean survival curve (left) and cell death time 50 (T_50_) (right) in human and naked mole-rat fibroblasts from skin and kidney of 60- and 30-days old animals. Human data (grey curve and bar) are the same as in Fig 1c. (d-e) Each dot (n) is the number of experiments. Human, mouse and naked mole-rat n=3. Student t-test (c) and One-way ANOVA (e). *p < 0.05, **p < 0.01, ***p < 0.001, ****p < 0.0001. Data are presented as mean values ± s.e.m.

**Extended data Fig 2.**
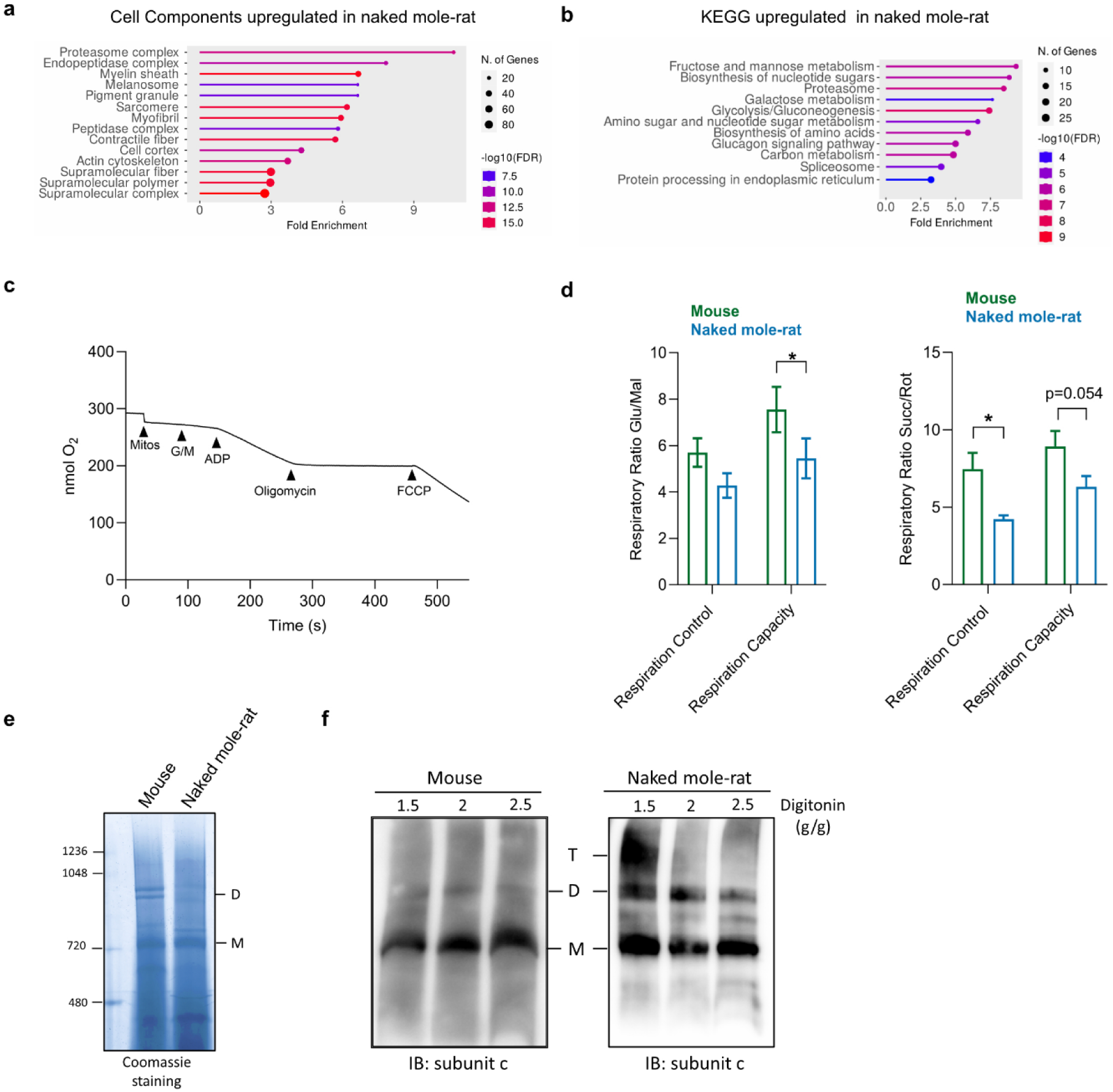
Comparison between mouse and naked mole-rat mitochondria. (a-b) Cell components and KEGG Gene Ontology (GO) term enrichment analysis of naked mole-rat upregulated proteins compared to mouse. **(c)** Representative trace of oxygen consumption measurements. The level of oxygen (nmol O_2_) is measured upon addition of mitochondria (Mitos), substrates (G/M), ADP, oligomycin, FCCP. **(d)** Respiratory control and respiratory capacity measured when either Glutamate/Malate (left) or Succinate/Rotenone (right) were provided. n is the number of animals, n=3. Multiple Student t-test. *p < 0.05. Data are presented as mean values ± s.e.m.**(e)** Blue native-PAGE analysis and subsequent Coomassie staining of mouse and naked mole-rat liver mitochondria. **(f)** Blue native-PAGE and subsequent immunoblotting against ATP synthase subunit c of mouse and naked mole-rat liver mitochondria in the presence of indicated amount of digitonin (g digitonin/g protein). Images are representative of two independent blots. M: monomers; D: dimers: T: tetramers.

**Extended data Fig 3.**
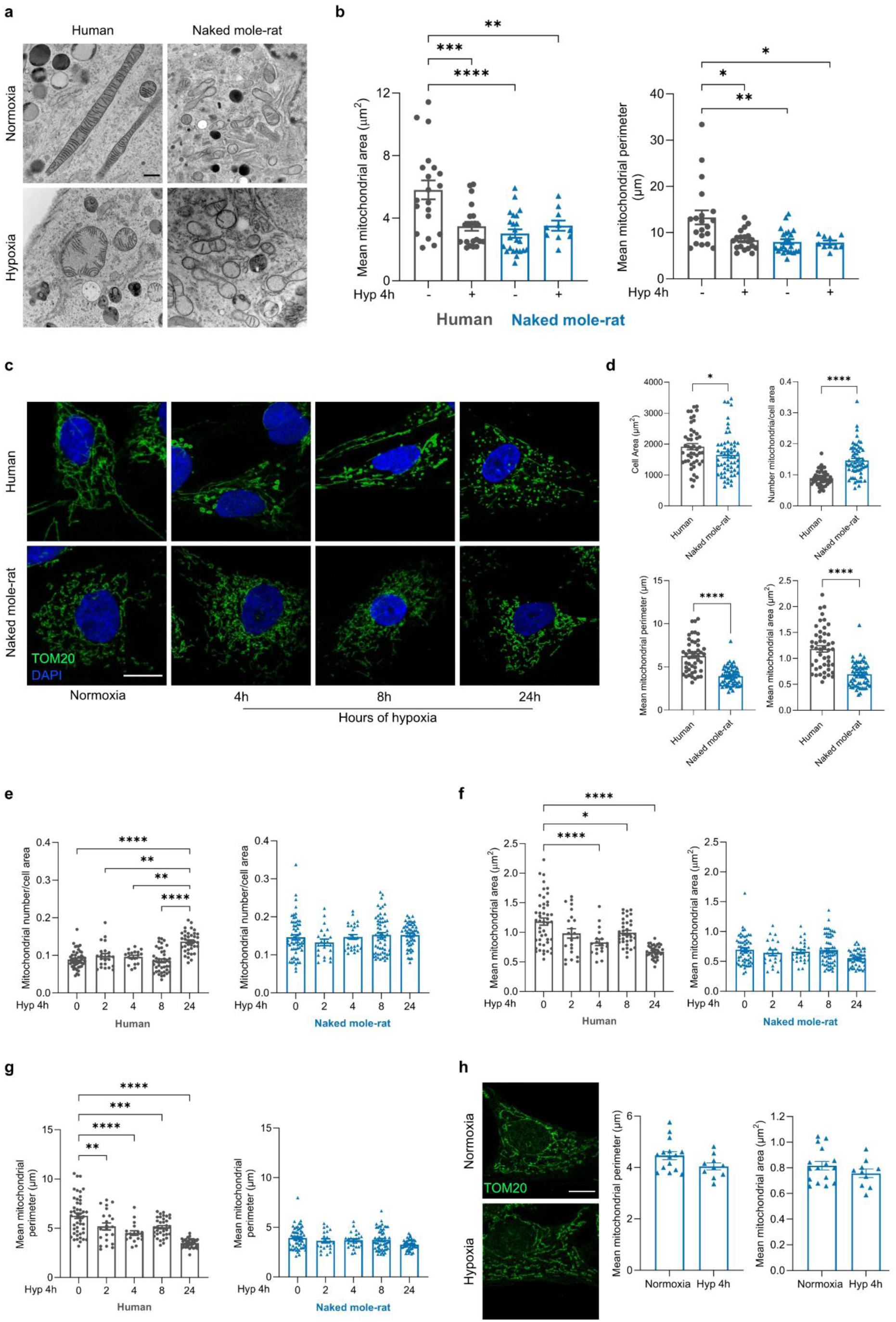
Naked mole-rats do not remodel their mitochondria up to 24 hours of hypoxia. (a-b) Transmission Electron Microscopy analysis of human and naked mole-rat mitochondria in normoxia and after 4 h of hypoxia (1%O_2_). (**a**) Representative pictures and (**b**) quantification of mitochondrial number, mean mitochondrial perimeter and area. Scale bar 500nm. Each dot (n) is number of cells, n>10 from 3 independent experiments. One-way ANOVA. **(c-d-e-f-g)** Mitochondrial morphology analysis in human and naked mole-rat fibroblasts in normoxia and after 2, 4, 8 and 24 h of hypoxia (1%O_2_). (**c**) Representative pictures, TOM20 (mitochondria, green), Dapi (nuclei, blue). Scale bar 10µm (**d**) Quantification of cell area (µm^2^) (top left), mitochondrial number/cell area (top right), mean mitochondrial perimeter (bottom left) and area (bottom right) in normoxic conditions in human and naked mole-rat. **(e)** Quantification of number of mitochondria/cell area, mean mitochondrial area (**f**) and perimeter (**g**) in human and naked mole-rat cells upon 2, 4, 8, 24 h of hypoxia. (**d-e-f-g**) Each dot (n) is number of cells, n>30 from 3 independent experiments. Normality test followed by Mann-Whitney test (d) and One-way ANOVA (e-f-g). Human and naked mole-rat data in normoxia are the same in d and in e, f, g. **(h)** Mitochondrial morphology analysis in naked mole-rat skin fibroblasts in normoxia and after 4 h of hypoxia (<1%O_2_). Right, representative pictures, TOM20 (mitochondria, green). Scale bar 10µm. Left, quantification of mean mitochondrial area and perimeter. Each dot (n) is number of cells, n>10 from 3 independent experiments. Student’s t test. *p < 0.05, **p < 0.01, ***p < 0.001, ****p < 0.0001. Data are presented as mean values ± s.e.m.

**Extended data Fig 4.**
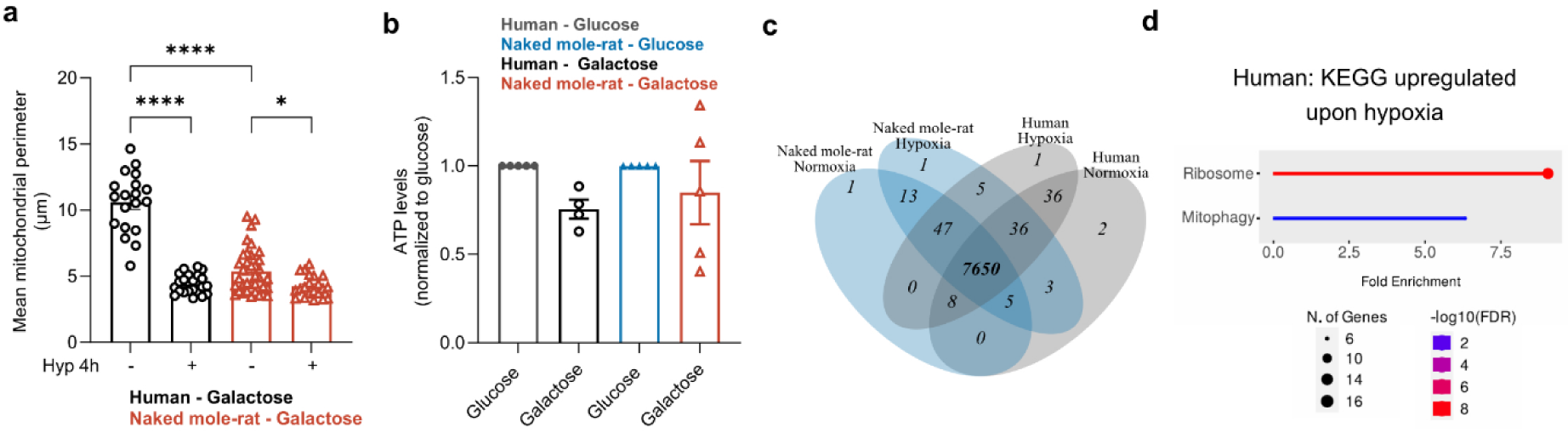
Oxygen deprivation affects mitochondrial functionality. **(a)** Quantification of the mean mitochondrial perimeter in human and naked mole-rat fibroblasts grown in “galactose medium” in normoxia and after 4 h of hypoxia (1%O_2_). n is number of cells; n>15 from 3 independent experiments. One-way ANOVA. **(b)** Total cellular ATP levels in human and naked mole-rat cells in glucose and galactose medium. Data were normalized to human or naked mole-rat cells in glucose medium. n is the number of experiments, n>3. One sample t-test **(c)** Number of proteins identified in the proteomic of human and naked mole-rat cells in normoxic and hypoxic (1%O_2_) conditions. **(d)** KEGG Gene Ontology (GO) term enrichment analysis of human upregulated proteins upon hypoxia. *p < 0.05, ****p < 0.0001. Data are presented as mean values ± s.e.m.

**Extended data Fig 5.**
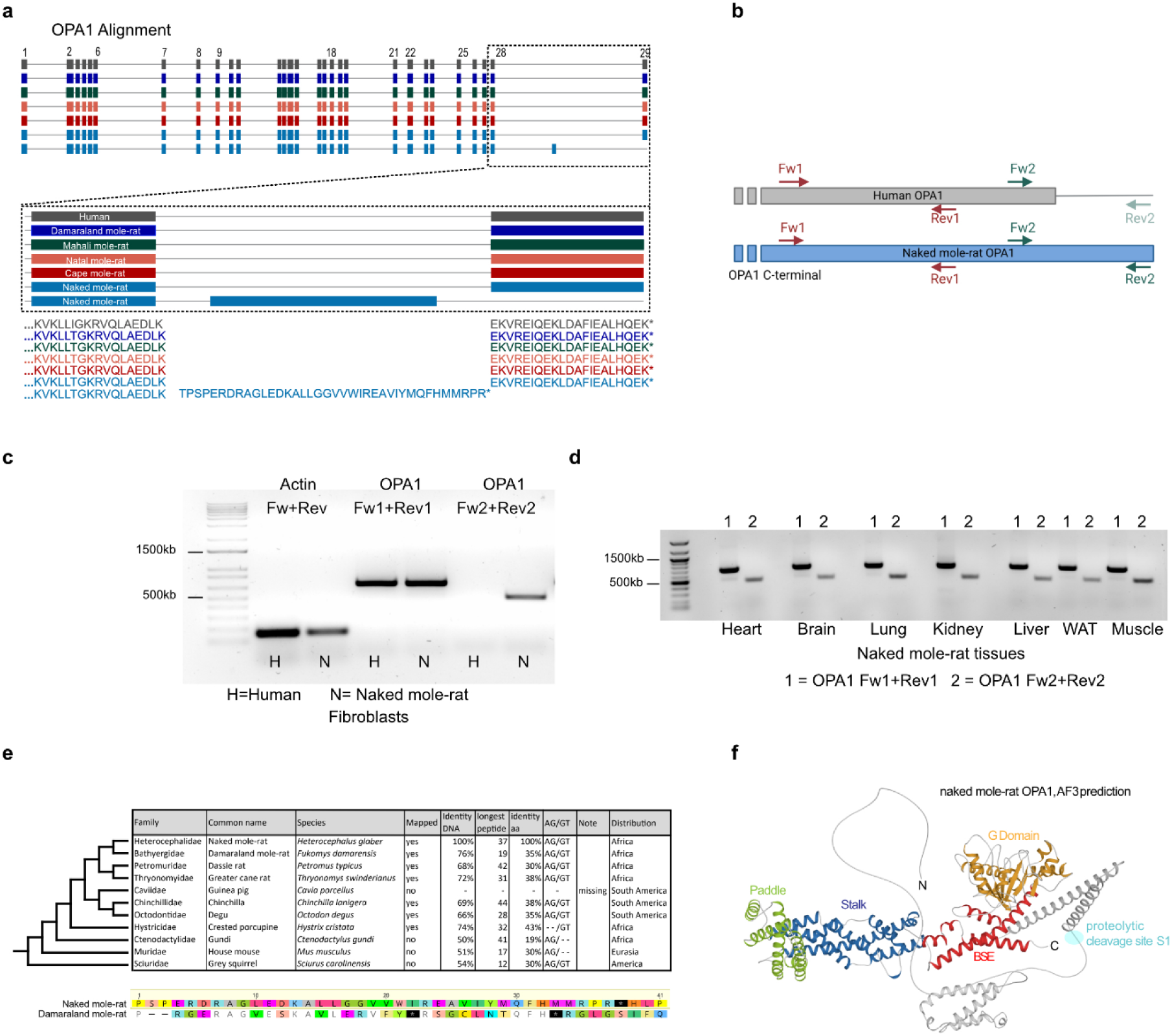
The naked mole-rat OPA1 contains a unique exon. **(a)** Transcriptomic sequence alignments of human and African mole-rat species OPA1. **(b)** RT-PCR primers scheme: Fw1, Rev1 and Fw2 are aligning to sequence conserved in human and naked mole-rat. Rev2 is aligning to the unique naked mole-rat C-terminal sequence. Actin primers were used as control. **(c-d)** RT-PCR performed on cDNA from human and naked mole-rat fibroblasts and from indicated naked mole-rat tissues. The resulting PCR product was purified and sequenced. **(e)** Comparison of the unique DNA exon sequence between naked mole-rat and the indicated species. Bottom, example of the DNA alignment between the naked mole-rat OPA1 unique exon sequence and the corresponding sequence from the Damaraland mole-rat. **(f)** Cartoon presentation of the predicted full length naked mole-rat OPA1. Color code for naked mole-rat OPA1 is as in Fig5B. The S1 proteolytic cleavage site is indicated in cyan.

**Extended data Fig 6.**
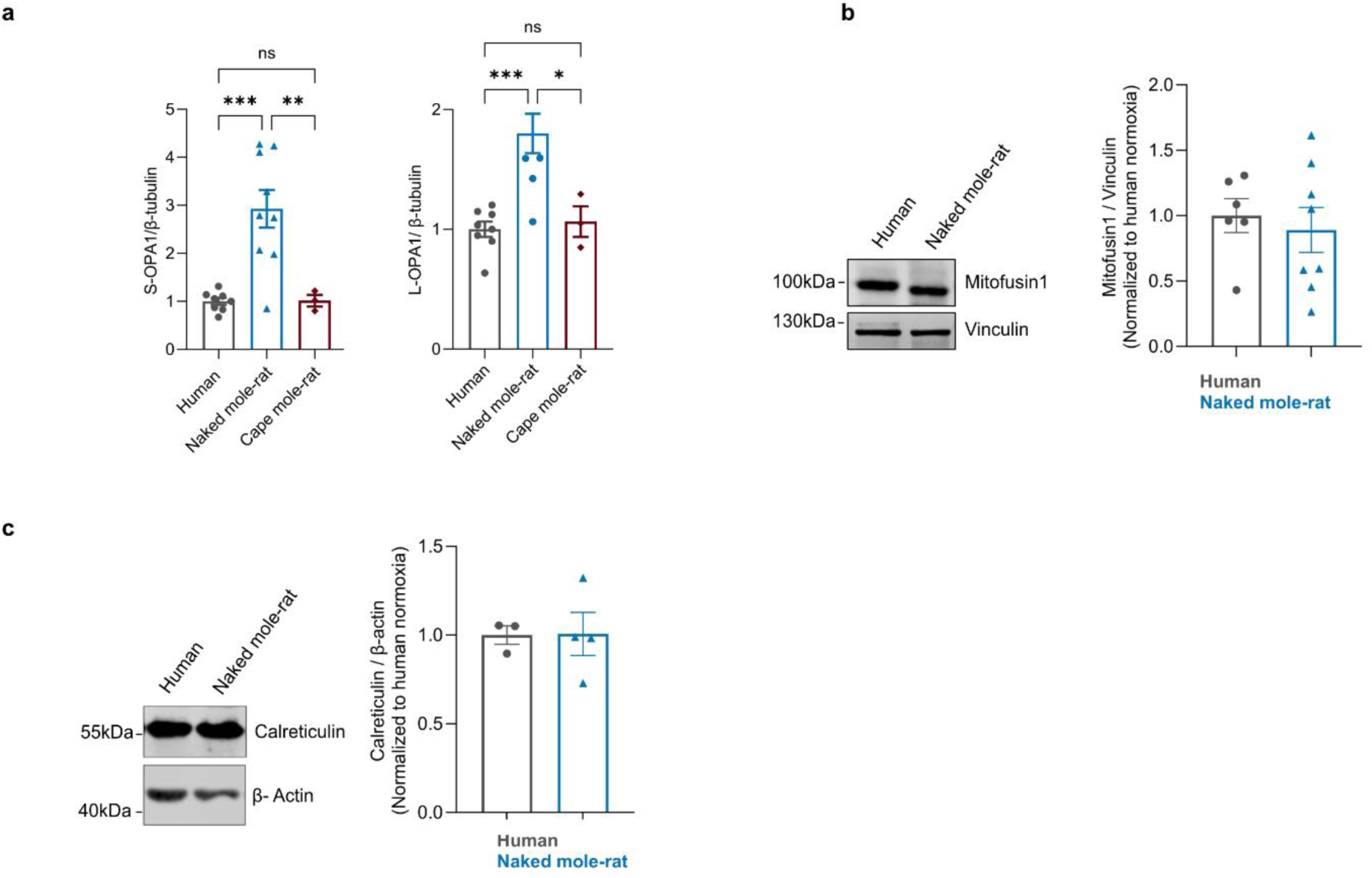
OPA1 levels in naked mole-rat, human and cape mole-rat. **(a)** Quantification of L-and S-OPA1 levels in human, naked mole-rat and cape mole-rat normalized to β-tubulin. Data are normalized to human. n is the number of experiments, n≥3. **(b)** Representative western blot (left) and quantification of Mitofusin levels in human and naked mole-rat cells normalized to Vinculin. Data are normalized to human. n is the number of experiments, n≥3. **(c)** Representative western blot (left) and quantification of Calreticulin levels in human and naked mole-rat cells normalized to β-Actin. Data are normalized to human. n is the number of experiments, n≥3. One-way ANOVA *p < 0.05, **p < 0.01, ***p < 0.001. Data are presented as mean values ± s.e.m.

